# Homeodomain transcription factors in adipocyte thermogenesis: insights into the species-specific and conserved regulatory elements of human UCP1

**DOI:** 10.1101/2024.05.16.594487

**Authors:** Beáta B. Tóth, Géza Hegedűs, Eszter Virág, Gergely Tamás Pethő, Levente Laczkó, László Fésüs

## Abstract

Adipocyte thermogenesis is a promising target for treating metabolic disorders, but its regulatory mechanisms remain unclear. This study investigates transcription factors (TFs) and regulatory elements that may control the human *UCP1* gene, which is essential for thermogenesis and the formation of the adipocyte phenotype. Using the Eukaryota Promoter Database, we performed computational analyses of the *UCP1, UCP2* and *UCP3* promoter sequences in humans, mice and rats to identify conserved and species-specific elements. We also used transcriptome data from human neck-derived adipocytes and databases (Contra v3, ChiPBase, TFlink, AdipoNet, ISMARA, etc.) to narrow potential regulatory TFs. Our results show that mouse and rat *UCP1* enhancers lack large segments, primarily due to the insertion of repetitive elements that are already lost in some clades. We identified key TFs such as PPARA, PPARG, THR, RARE, RXR, JUN, TFAP2 and SREBF1 as general regulators of *UCPs.* Additionally, human-specific *UCP1* regulatory hotspots (e.g. 5’-TCTAATTAGA-3’) recognised by homeodomain TFs (e.g. EN1, PAX4, HOXA5 and PRRX2) and NFIL3 were detected. Phylogenetically conserved regulatory elements suggest common TFs in human *UCP1* paralogues (MAX, MYCN, MNT, HES1) and cross-species (POU6F1). These results improve our understanding of thermogenic adipocyte development and provide new therapeutic targets for metabolic diseases.

**Graphical Abstract:** 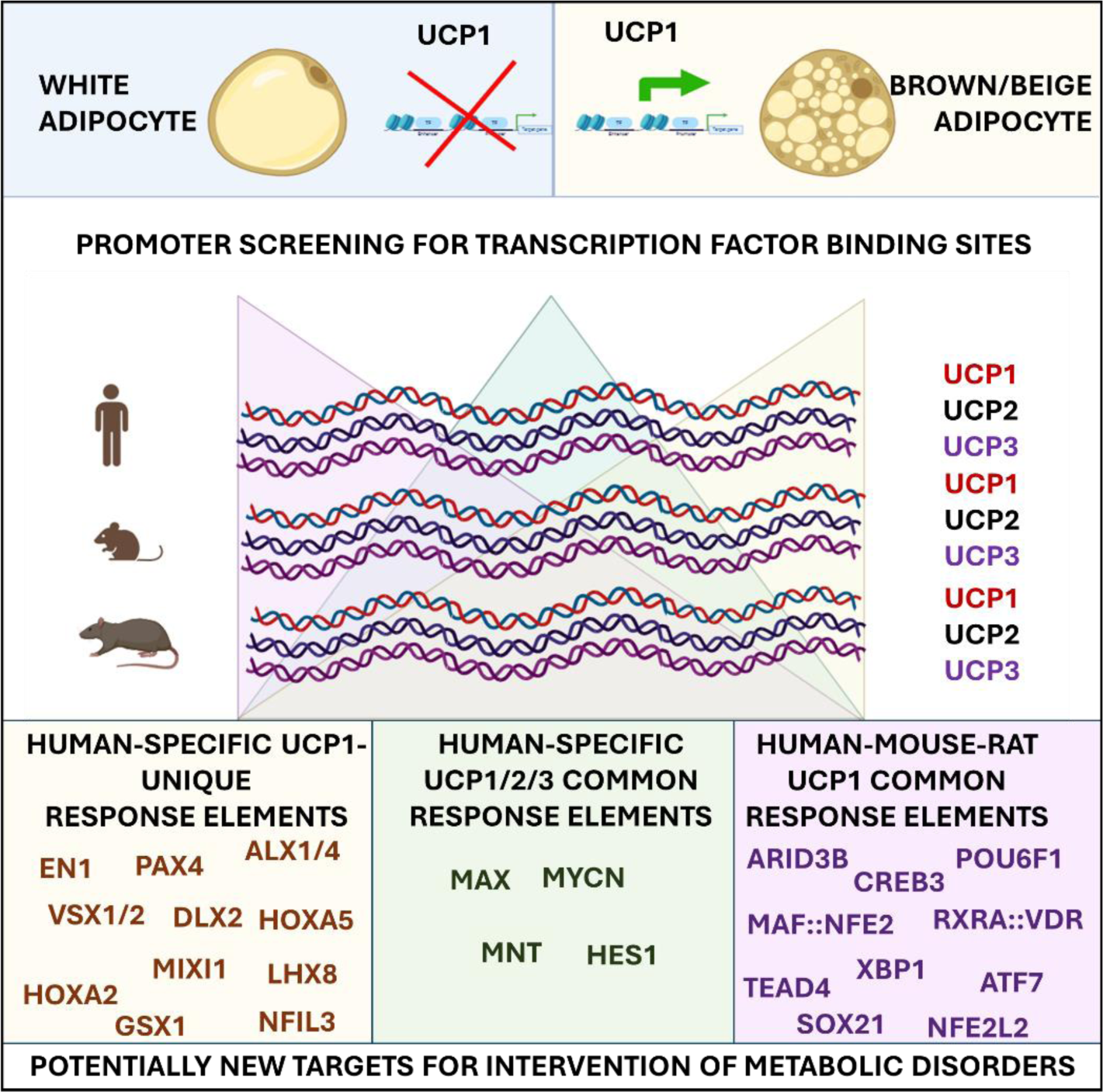

## I. INTRODUCTION

Adipocytes are crucial for maintaining energy balance and a healthy metabolism in the human body. While white adipocytes primarily serve as energy reservoirs, brown and beige adipocytes are specialised in energy expenditure through thermogenesis. This thermogenic capacity is promising for influencing overall energy homeostasis, especially in the context of metabolic disorders (Tayanloo-Beik et al., 2023; Seki et al., 2022; Wibmer et al., 2021; Becher et al., 2021; Razzoli et al., 2016, Nedergaard et al., 2007). Non-shivering adipocyte thermogenesis, a crucial component of the body’s heat production mechanisms, can be triggered by cold exposure and various environmental stimuli, however, the intricate mechanisms are not yet fully understood (Hoang et al., 2013). Cold-induced adaptive thermogenesis involves the upregulation of Uncoupling Protein 1 (UCP1) in the mitochondrial membrane, which recycles protons to the matrix, thereby uncoupling oxidative phosphorylation and ATP synthesis. This leads to controlled heat production and is an attractive target for weight control strategies (Feldman et al., 2009; Ricquier 2011, Nedergaard et al. 2001, Enerbäck 1997). In addition to UCP1-mediated thermogenesis, other cyclic mechanisms may also contribute to adipocyte thermoregulation, such as endoplasmic reticulum-associated Ca2+ (Ikeda et al. 2017), mitochondrial creatine (Kazak and Spiegelman 2020) and the triacylglycerol-fatty acid cycle (Cohen and Kajimura 2021). Despite extensive research, a comprehensive understanding of the regulatory mechanisms controlling adipocyte thermogenesis/browning remains elusive (Waldén et al. 2012; Nedergaard and Cannon 2014). Several TFs have been proposed as direct regulators of *UCP1* gene expression, including peroxisome proliferator-activated receptors (PPARs), retinoic acid receptors (RXRs, RARs), thyroid hormone receptor (THR), cAMP response element-binding proteins (CREBPs), nuclear factor erythroid response element 2 like 2 (NFE2L2), sterol regulatory element-binding protein 1 (SREBP1), early B-cell factor 1-2 (EBF1-2), the c-Jun::c-Fos complex and the CCAAT/enhancer- binding proteins (CEBPA) as well as indirect regulators such as PPARG coactivator 1 alpha (PGC-1α) and PR domain containing 16 (PRDM16) (Rosen et al. 2000; del Mar Gonzalez-Barroso 2000; Rim and Kozak 2002; Seale et al. 2007; Timmons et al. 2007; Ricquier 2011; Villaroya et al. 2017; Zhang et al. 2023). Recent reviews provide insights into the complex epigenetic and transcriptional regulation underlying brown/beige adipocyte development (e.g. Li et al. 2019; Shapira and Seale, 2019; Ambele and Pepper, 2017; Villaroya et al., 2017; Inagaki et al., 2016; Harms et al., 2015; Bonet et al., 2013), but the intricate web of regulatory mechanisms is still to some extent unclear.

This study aims to use computational methods to identify transcription factors (TFs) that may regulate the expression of the human *UCP1* gene by identifying TFs that have putative functional DNA-binding motifs within the promoter regions of the human UCP1 gene in the range of -5Kb – +1Kb (6000 base pairs (BP); position: GRCh37/hg19; Chr. 4: 141,489,115- 141,495,114). To simplify nomenclature, the term ’promoter’ is used here for the entire sequence regions analysed, including the intergenic regions (1kb DNA segments in the positive direction from the transcription start site (TSS)), the proximal promoter (-1kb – TSS) and the distal enhancers (-1kb – - 5kb). By comparing *UCP1* orthologues and paralogues in humans, mice and rats, we aimed to uncover conserved and human-specific *UCP1* regulatory elements.

To determine whether a particular binding sequence is present within the human *UCP1* promoter, we used the eukaryotic promoter database (EPD) interface. This database uses a probability-based algorithm to identify TF response elements (REs) at four different p-value thresholds (p-value < 0.01, 0.001, 0.0001 and 0.00001). In this study, we focused on the two most stringent p-value thresholds (p < 0.0001 and p < 0.00001) because empirical evidence and computational results suggest that more permissive thresholds may lead to high false-positive rates (Wunderlich and Mirny, 2009; Benita et al., 2009). However, our exploratory studies have shown that using the most stringent threshold (p < 1e-5) can lead to high false-negative rates, as in the case of RE of HIF1A TF in gene promoters such as *VEGFA* and *GLUT1*, where the regulatory role of HIF1A is well known. By setting the significance level at p < 1e-5, there would certainly be fewer false positives in the data, but the number of false negatives would increase and be lost for further analysis, while false positives can be filtered out with high efficiency by comparing paralogous and orthologous gene promoters. Therefore, to achieve a balance between sensitivity and specificity, we primarily used the threshold p < 1e- 4, while occasionally referring to the results of p < 1e-5.

To exclude false results, we extended our analysis to *UCP2* and *UCP3* promoters from humans, mice and rats. The UCP proteins belong to the solute carrier family (SLC), which consists of different transporters such as ion channels, exchangers and passive transporters. *UCP2* and *UCP3* are paralogues of *UCP1* but have different tissue distributions and do not share its thermogenic function (Nedergaard and Canon 2003, Rousset et al. 2004, Nicholls 2021). Both UCP2 and UCP3 contribute to the fine-tuning of mitochondrial function and energy metabolism and are therefore important players in the maintenance of metabolic health and protection against metabolic disorders, but their exact function is still unclear. UCP2 is widely distributed in various tissues, including adipose tissue, brain, lung, pancreas and immune cells. In contrast to UCP1, which is mainly found in brown adipose tissue and is primarily involved in non-shivering thermogenesis by conducting protons back into the matrix, UCP2 has a more general role. UCP2 is thought to regulate the production of reactive oxygen species (ROS) and thus protect cells from oxidative stress (Hirschenson et al. 2022). UCP2 is also involved in the regulation of insulin secretion by beta cells and in this way can also influence metabolic efficiency and energy expenditure (Cadenas 2018). UCP3, on the other hand, is mainly expressed in skeletal muscle and to a lesser extent in brown adipose tissue. One of the main proposed functions of UCP3 is to promote the export of fatty acids from the mitochondria, thereby preventing lipid-induced oxidative damage (Himms-Hagen and Harper 2001, Schrauven et al. 2002). UCP3 may also be involved in maintaining energy balance and may play a role in muscle thermogenesis, especially during periods of increased fatty acid intake or muscle activity. By comparing these closely related genes, we aimed to identify TFs specifically involved in the regulation of UCP1 and thereby adipocyte thermogenesis.

We also aimed to identify conserved and human-specific regulatory elements of the *UCP* promoters by studying evolutionarily distant orthologues from mouse and rat. While mice and rats are valuable as model organisms for thermogenic studies, the translation of results to human physiology can be challenging due to inherent differences (Enerbäck et al. 1997). These differences may affect metabolism and thermogenic processes, as variations in body size and environmental temperature are among the factors that play an important role in shaping evolutionary strategies to maintain energy balance (Tattersall and Milsom, 2009; Kottke et al., 1948). Exploring potential regulatory differences in the expression of paralogous genes with common origins but different tissue expression and protein functions, such as *UCP2* and *UCP3*, in various species opens up the possibility of deciphering human *UCP1*-specific regulators and providing insights into potentially more specific targets of intervention that may not be apparent in traditional model organisms (Nedergaard and Canon 2003, Rousset et al. 2004, Nicholls 2021, Hughes and Criscuolo 2008, Enerbäck et al. 1997, Warfel et al. 2023).

This study aims to provide a comprehensive understanding of the transcriptional regulation of human *UCP1* by identifying key TFs with putative binding motifs. We take advantage of functional differences between members of the UCP family and evolutionary conservation between species. This could elucidate the specific regulatory processes that control the phenotype of human adipocytes and improve our understanding of thermogenic fat cell development, which would have significant therapeutic implications.

## RESULTS

### 1. Comparative analysis of the sequences of *UCP1*, *UCP2* and *UCP3* promoters, genes and proteins in different species: structural and evolutionary aspects

To elucidate the structural features underlying the specific function of the three mitochondrial solute carrier family 25 (SLC25) proteins, UCP1 (SLC25A7), UCP2 (SLC25A8) and UCP3 (SLC25A9), we performed a thorough comparative analysis encompassing the nucleotide sequences of the genes and their promoters in the human, mouse and rat genomes. We also aligned the amino acid sequences of the encoded proteins and analysed their complex three-dimensional (3D) spatial structures.

#### 1.1. Sequence similarity analysis shows distinct evolutionary trends between genes and their regulatory regions

A phylogenetic tree reconstructed by merging the human, mouse and rat *UCP1, UCP2* and *UCP3* data clearly shows that orthologous sequences were grouped, indicating a closer relationship than to their paralogous counterparts when considering amino acid composition, coding (mRNA) and regulatory DNA sequences, with high support values in all cases (Fig. 1A). As expected, orthologous genes also show greater similarity between mouse and rat and are more tightly clustered compared to human. In contrast, we obtain a different clustering of the paralogous genes depending on the sequences analysed. While the reconstruction of the mRNA and amino acid sequences identified UCP2 and UCP3 as a separate cluster from UCP1, UCP1 and UCP2 were grouped based on the promoter sequence. These results are also confirmed by pairwise distance analysis of the coding and promoter sequences of the *UCP* genes (Suppl Fig. 1A).

**Figure 1.**
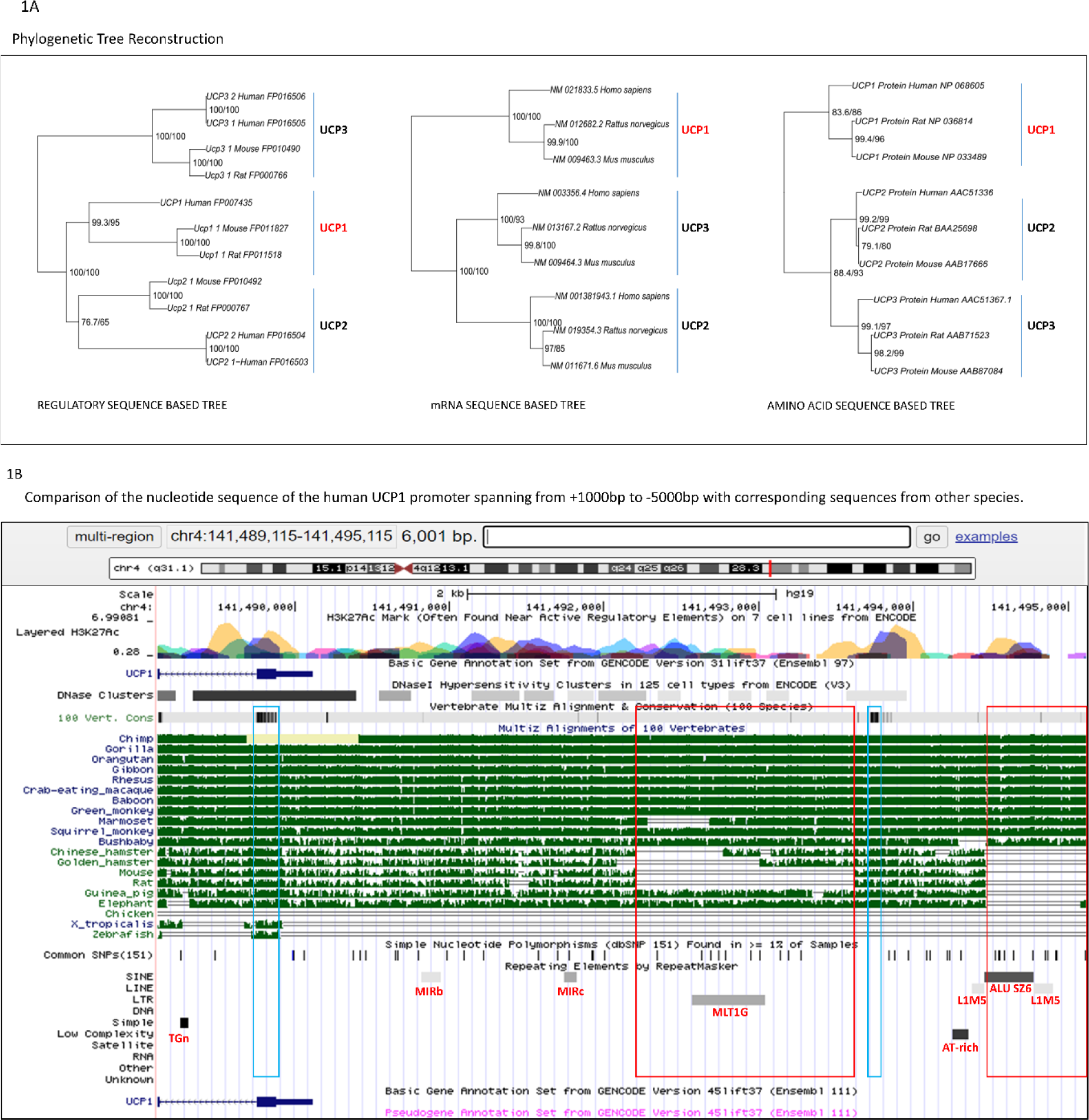
**1A.** Phylogenetic tree reconstruction shown as a midpoint rooted phylogram based on the nucleotide sequence of the *UCP1*, *UCP2* and *UCP3* upstream regulatory region, the mRNA and amino acid sequence of the corresponding proteins in human, mouse and rat. Support values (aLRT/rapid bootstrap) are shown above the branches (packages ape 5.7.1 in R environment). **1B** Comparison of the nucleotide sequence of the human UCP1 promoter between +1000bp – -5000bp with sequences from other species using the Multiz Alignment of 100 Vertebrates database in the UCSC genome browser. Red boxes highlight sequence segments absent from *UCP1* promoter of mice and rats, blue boxes indicate segments displaying significant conservation across diverse species. Red letters and grey lines indicate the abbreviation and position of the repetitive elements detected by UCSC integrated Repeat Masker database.

#### 1.2 The presence of transposable and repetitive elements leads to considerable sequence divergence within the distal enhancer regions of humans and rodents

The thermogenic *UCP1* gene evolved around 120-150 million years ago after the split of the platypuses (monotremes) and marsupials (metatherians) from the lineage that gave rise to the placental mammals (eutherians) (Gaudry et al. 2017). However, Jastroch et al. (2005) demonstrated that all three members of the core *UCP* family were present before the divergence of the ray-finned and lobe-finned vertebrate lineages around 420 million years ago, however, were later lost or inactivated in several clades, such as birds. A cross-species alignment of the analysed 6000 bp long nucleotide sequences of the *UCP1* promoter revealed that a 1419 bp long sequence interval extending from -2083 bp to -3502 bp (GRCh37/hg19: Chr4 141.492.198-141.493.617) in the distal enhancer region of the human *UCP1* promoter and the last 658 bp long segment of the 6000 bp long sequence analysed (Chr4: 141.494.458- 141.495.115; at -4341 bp to -5000 bp) was missing in the UCP1 promoters of mice and rats (Fig. 1B). This absence was confirmed by a BLAST search in the NCBI database (https://blast.ncbi.nlm.nih.gov/). Examination of the presence of the 1419 bp-long sequence in other species using the Multiz Alignment of 100 Vertebrate Tool in the UCSC Genome Browser (https://genome.ucsc.edu/) suggests that it is an evolutionarily novel sequence that arose around the divergence between placentals and marsupials with the appearance of the thermogenic *UCP1* gene (Jastroch et. al. 2024), however, it seems to be secondarily lost in certain clades, including mice and rats (Suppl. Fig. 1B). It is noteworthy that short palindromic sequences or their residues can be detected at the beginning and end of these evolutionarily novel DNA fragments in the human *UCP1* promoter (Suppl. Fig. 1C, 1D,1E, 1F, 1G; red boxes). Eukaryotic Promoter Database analysis (p < 1e-5) shows that the RE of CREB3 at -3498 bp corresponds to the palindromic “TGACGTCA” sequence at the end of the 1419 bp long segment (Suppl. Fig. 1D). Notably, the 468 bp central region of this new sequence (Chr4: 141.492.573-141.493.040; at -2458 bp to -2925 bp), identified as a long terminal repeat (LTR) retrotransposon repetitive element by RepeatMasker (http://www.repeatmasker.org) (Jurka 2000) named MLT1G from the endogenous retroviruses ERVL-MaLR family (Fig. 1B). This 468 bp sequence appears to be integrated into the human genome several times, as the BLAST search shows (Suppl Fig. 1H). It is also interesting that at the beginning and the end of this central segment, TF REs of the homeodomain family are identified, DLX2 RE at -2478 bp, while at the end for HOXD8, HOXD9, HOXB5, POU6F1 at -2913 bp. The Eukaryotic Promoter Database does not identify a TF RE in the middle part of this central segment with a probability of p < 1e-5. In the 658 bp long sequence at the end of the UCP1 promoter segment which was also missing in the rodent genome, a 317 bp (Chr4: 141.494.458-141.495.774; at -4343 bp to -4659 bp) and a 124 bp long segment (Chr4: 141.494.775-141.494.898; at -4660 bp to -4783 bp) were also recognised by RepeatMasker as a short interspersed nuclear element (SINE) and a long interspersed nuclear element (LINE), respectively (Fig. 1B). The SINE was classified as an Alu sz6 repeat from the ALU family, and the following LINE was identified as an L1M5 repeat which belongs to the L1 family. The BLAST searches have found similar repeat sequences at over 1000 sites in the human genome. Using RepeatMasker, we saw several additional repetitive elements in the human *UCP1* promoter, whose possible role requires further investigation (Fig. 1B, grey lines). This observation may emphasise the dynamic nature of the genetic elements and their potential role in regulating gene expression.

Nevertheless, the intronic regions of *UCP1* at around +250 bp and the distal enhancer region at around -3600 bp show above-average sequence similarity across species and may indicate conserved DNA regions with REs that are important for transcriptional control, such as the myeloid zinc finger 1 (MZF1) at -3667 bp, and the glucocorticoid modulatory element binding protein 1 (GMEB1) and GLIS family zinc finger 2 (GLIS2) at around +250 bp (Fig. 1B, blue boxes; Suppl Table 2).

#### 1.3 Subtle structural differences between orthologous and paralogous UCP1 proteins may have functional implications

The primary 3D structures of human *UCP1, UCP2* and *UCP3* proteins show a high degree of homology with their mouse and rat counterparts. This was established by predictive modelling with AlphaFold 2 and structural alignment with the RCSB PDB interface (Fig. 2A, see Methods). Supplementary Figure 2A shows these spatial protein structures in a colour-coded format and provides insight into their conformational similarities. Remarkably, the model reliably predicts the spatial structure of *UCP1* in all three species, whereas model confidence tends to be lower for *UCP2* and *UCP3*. Interestingly, a short interdomain sequence shows a particularly low model confidence within the UCP proteins of all three species (Suppl Fig. 2A, red colour). The context-dependent variability in spatial structure may provide the opportunity to influence functionality under certain conditions, such as modulating proton conductance or influencing the binding of fatty acids and purine nucleotides. Detailed pairwise structural alignment measurements of the analysed proteins, showing the extent of structural similarity to human *UCP1*, are presented in Table 1. These analyses show a consistent template modelling score (TM score) of 0.75 or higher compared to human UCP1 and matching equivalent residue numbers in all cases (Table 1). In particular, the results show the remarkable similarity between mouse and rat *UCP1* and human *UCP1* with an impressive TM score of 0.97. The homology is slightly lower for the paralogous genes *UCP2* (0.84 TM score) and *UCP3* (0.77 TM score). It is worth noting that the amino acid sequence similarities closely match the result of the DNA sequence comparisons (Table 1.), both underscoring the greater similarity in the orthologous protein structures. However, the relevance of amino acid sequence homology to protein structure similarity is challenged by the fact that the presence of the conserved UCP protein structure is a recurring feature within the SLC25 protein family (Hoang et al., 2013; Crichton et al., 2017), regardless of the degree of homology between the constituent amino acids. For example, the ADP/ATP translocase (SLC25A4 or ANT1) has a similar predicted 3D folded structure (Fig. 2B) and a substantial TM score of 0.8, despite a modest amino acid homology of 21% (Table 1). In summary, the phylogenetic analyses of *UCP1, UCP2* and *UCP3* in the human, mouse and rat genomes, and their protein 3D structural comparison show the closest relationships between the orthologues. The paralogues show different clustering patterns based on gene and promoter sequences. The observed structural similarities, independent of amino acid sequence homology, emphasise the conserved nature of UCP protein structure within the SLC25 family and raise questions about the functional impact of sequence variations in specific interdomain regions.

**Figure 2.**
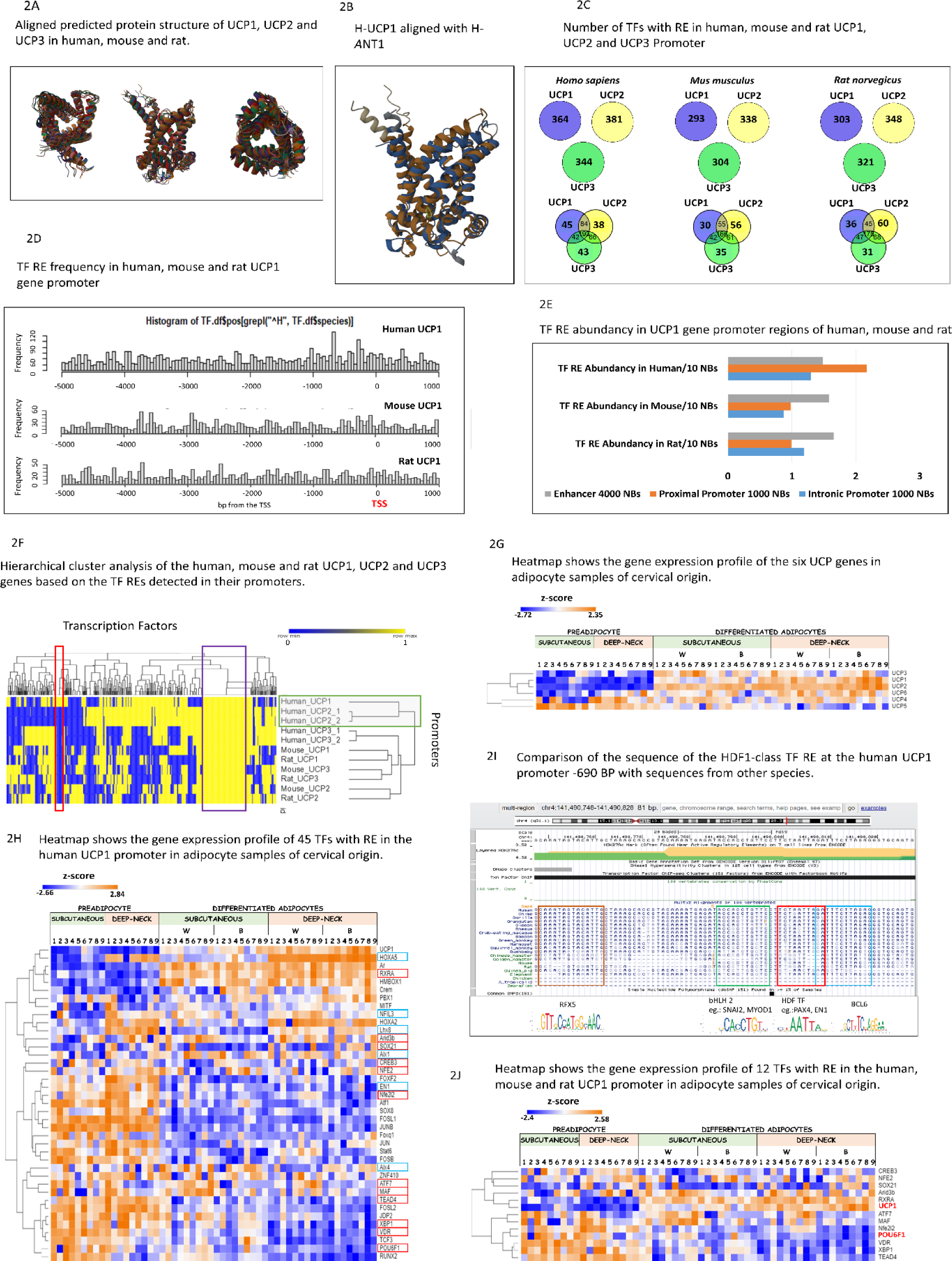
**1A**. RCSB PBD aligned, AlphaFold 2 predicted protein structures of human, mouse and rat UCP1, UCP2 and UCP3 from three different perspectives. **1B.** AlphaFold 2 predicted H-UCP1 aligned with H-ANT1. **1C.** Number of TFs with RE in human, mouse and rat *UCP1*, *UCP2* and *UCP3* Promoter: -5000 bp – +1000 bp (p<1e-4) (Venn program in R environment). **1D.** TF RE frequency in human, mouse and rat *UCP1* gene promoters -5000 bp – +1000 bp (p<1e-4) (Excel). **1F.** Bars show the frequency of TF RE per 10 bp in different parts of the UCP1 promoter in humans, mouse and rat -5000 bp – +1000 bp (p<1e-4). Distal enhancer -5000 bp – -1001 bp; Proximal and Core promoter-1000 bp – TSS; Intronic promoter TSS – +1000 bp (Excel). **2F**. Hierarchical cluster analysis of the human, mouse and rat *UCP1*, *UCP2* and *UCP3* genes based on the TFs detected in their promoters by RE. The red box highlights the 13 TFs that are unique to the human *UCP1* promoter, the purple box shows the 93 TFs that are common to the examined UCPs, while the green box shows the clusters with the most similar TF RE profile to *UCP1* (Morpheus Interface). **2G** Heatmap shows the gene expression profile of the six *UCP* genes in adipocyte samples of cervical origin. Hierarchical cluster analysis based on the Spearman Rank similarity matrix identifies *UCP*s with similar gene expression profiles to *UCP1* (Morpheus Interface). **2H** Heatmap shows the gene expression profile of 45 TFs with RE in the human *UCP1* promoter in adipocyte samples of cervical origin. For comparison, *UCP1* is also shown. Hierarchical cluster analysis based on the Spearman Rank similarity matrix identifies TFs with similar gene expression profiles to *UCP1* (Morpheus Interface). **2I** Comparison of the nucleotide sequence of the HDF1-class TF RE at the human *UCP1* promoter -690 bp with sequences from other species using the Multiz Alignment of 100 Vertebrates database in the UCSC genome browser (red box). In the vicinity of HDF1 RE the TF RE sequences of RFX5, MYOD1 and BCL6 identified by EPD are highlighted with brown, green and blue boxes, respectively. **2J** Heatmap shows the gene expression profile of 12 TFs with RE common in the human, mouse and rat *UCP1* promoter in adipocyte samples of cervical origin. For comparison, *UCP1* is also shown. Hierarchical cluster analysis based on the Spearman Rank similarity matrix identifies TFs with similar gene expression profiles to *UCP1* (Morpheus Interface). Adipocytes were differentiated using a white (W) or brown (B) differentiation protocol (Tóth et al. 2020).

**Table 1.** Results of pairwise structure alignments, measures describing the extent of structural similarity of the human, mouse and rat UCP1, UCP2 and UCP3 proteins to human UCP1. The structure alignments are based on protein structures predicted by AlphaFold 2 and aligned by RCSP-PBD. For interpretation of the measures, see Methodology.

|  | Entry | Chain | RMSD | TM-score | Identity | Equivalent Residues | Sequence Length | Modelled Residues |
| --- | --- | --- | --- | --- | --- | --- | --- | --- |
| UCP1-Human | AF-P25874-F1-model_v4.pdb | A | - | - | - | - | 307 | 307 |
| UCP1-Mouse | AF-P12242-F1-model_v4.pdb | A | 1.14 | **0.97** | 79% | 307 | 307 | 307 |
| UCP1-Rat | AF-P04633-F1-model_v4.pdb | A | 1.06 | **0.97** | 79% | 305 | 307 | 307 |
| UCP2-Human | AF-P55851-F1-model_v4.pdb | A | 3.08 | **0.84** | 57% | 304 | 309 | 309 |
| UCP2-Mouse | AF-Q9R246-F1-model_v4.pdb | A | 2.86 | **0.84** | 57% | 301 | 309 | 309 |
| UCP2-Rat | AF-P56500-F1-model_v4.pdb | A | 2.2 | **0.89** | 57% | 302 | 309 | 309 |
| UCP3-Human | AF-P55916-F1-model_v4.pdb | A | 3.51 | **0.77** | 48% | 298 | 312 | 312 |
| UCP3-Mouse | AF-P56501-F1-model_v4.pdb | A | 4.01 | **0.75** | 54% | 305 | 308 | 308 |
| UCP3-Rat | AF-P56499-F1-model_v4.pdb | A | 3.55 | **0.78** | 46% | 299 | 308 | 308 |
| ANT1-Human | AF-P12235-F1-model_v4.pdb | A | 3.07 | **0.8** | **21%** | 283 | 298 | 298 |

### 2. Comparative analysis of transcription factor binding sites of *UCP* promoters in humans, mice and rats

#### 2.1 Cross-species comparative analysis of putative TF REs in the UCP1/2/3 promoters reveals greater similarity of human UCP1 to human UCP2 than to rodent UCP1

After a thorough investigation of the promoter sequences of the *UCP1*, *UCP2* and *UCP3* genes in humans, mice and rats, we focussed on determining the specific transcriptional regulation of the human *UCP1* gene. Using the Eukaryotic Promoter Database (EPD) and a large number of TF motifs from the JASPAR databases, we performed an in-depth analysis with 579 TF motifs (Dreos et al., 2017). Using stringent significance thresholds (p < 1e-4), we identified a variable number of TFs with putative REs in the nine genes analysed, ranging from 293 to 381 (Fig. 2C, Suppl Fig. 2B, Suppl Table 1). Applying a more stringent threshold of p < 1e-5 resulted in the detection of 63 to 123 TFs with REs in the promoter of the *UCP* genes in the three species (Suppl Fig. 2C). The distribution of TF REs within the *UCP1* promoters showed considerable differences between regions and species. While in humans the proximal promoter has the highest TF binding sites, in mice and rats it is the enhancer region (Fig. 2D, 2E; Suppl Table 2). Visualisation on a heatmap and hierarchical clustering showed that the regulatory TF-RE profile of the human *UCP1* gene is very similar to that of its paralogue, the human *UCP2* gene (Fig. 2F). This is in contrast to the observations based on protein structure, amino acid composition, and gene and promoter sequences (Fig. 1A, 2A), where the orthologues are more similar. Comparing the relative gene expression profiles of the human *UCP* genes in the cervical-derived adipocyte samples, the expression of *UCP1* was most closely related to that of *UCP2*, as shown by the correlation using Spearman’s Rank Similarity Matrix (Fig. 2E).

#### 2.2 Comparative analysis of transcription factor response elements of 9 UCPs reveals unique regulatory sequences for the human UCP1 gene associated with TFs of the homeodomain family

Our comprehensive study identified a total of 364 TFs (including dimers, trimers and variants) with putative REs in the analysed promoter region of the human *UCP1* gene (Figure 2C; Suppl Table 1). A comparison with the results of the analyses of the regulatory regions of the human *UCP2* and *UCP3* genes shows that 319 of these TF REs were also present in one of the analysed paralogous human *UCP1* promoters. However, 45 TFs (including dimers and variants) were present exclusively on the *UCP1* promoter. We examined these 45 *UCP1*-specific TFs in more detail by first comparing their expression patterns with *UCP1* in cervical-derived subcutaneous and deep neck preadipocytes and mature adipocytes, differentiated ex vivo (Figure 2H). Gene expression of 5 TFs, including *AR, RXRA, HMBOX1, CREM* and *HOXA5* showed a significant positive correlation with *UCP1* (Suppl Table 3). Conversely, the gene expression of 17 TFs, including *JUNB, FOSL1* and *2, ATF1, VDR* and *NFE2L2* as the top six, showed a negative correlation. However, the RE of RXRA shows only a specific occurrence in the human *UCP1* promoter when dimerised with VDR. Similarly, only a single variant of the JUNB TF RE, JUNB_var2 RE, shows a specific occurrence in the human *UCP1* promoter and only when dimerised with FOSL1 or FOSL2. When considered alone or dimerised with other TFs, their REs are identified in all *UCP*s tested, so their *UCP1* specificity in humans is context-dependent.

Next, in comparison to the TFs with REs detected in the regulatory regions of mouse and rat *UCP1, UCP2*, and *UCP3* genes, we identified 13 of the 45 human *UCP*1-specific TFs that do not have REs in the promoters of these rodent *UCP* genes, namely: ALX1, ALX4, DLX2, EN1, GSX1, HOXA2, LHX8, MIXL1, NFIL3, PAX4, VSX1, VSX2 and HOXA5 (Figure 2F, red box). Except for NFIL3, these TFs belong to the class of homeodomain family TFs (HDF; JASPAR database). The Eukaryotic Promoter Database can also detect most of these REs with the highest probability (p < 1e-5) (Table 2A, Suppl Table 2), supporting that these TFs may regulate thermogenic processes in a species-specific manner. In addition, we examined the RE nucleotide sequences identified by the Eukaryotic Promoter Database for these 13 TFs and detected a palindromic DNA nucleotide sequence of 10 bp, 5’-TCTAATTAGA-3’, positioned at -690 bp in the human *UCP1* promoter (EPD database; UCSC Genome Browser GRCh 37/hg19: 141.490.801- 141,490,810), and serves as RE for 9 of the 13 human *UCP1*-specific TFs (Table 2A). While this short RE sequence was located outside the 1419 bp and 658 bp long sequence region found only in humans, it was indeed absent from the mouse and rat *UCP1, UCP2*, and *UCP3* promoters, as well as the human *UCP2* and *UCP3* promoters, according to Eucaryotic Promoter Database and NCBI BLAST searches, suggesting that it potentially play a unique regulatory role in human adipocyte thermogenesis. Further data indicate that this sequence is conserved to some degree in primates (one base difference), such as chimpanzees, gorillas, orangutans, baboons and green monkeys (Figure 1B; Multiz Vertebrate Sequence Alignment in UCSC), expanding the range of possible species in which this regulatory process may be implemented. In addition to the homeodomain family TFs, the nuclear factor interleukin 3 (NFIL3), which belongs to the basic leucine zipper TF family and binds to the activating transcription factor (ATF) site, was found to be another potential human-specific *UCP1* regulatory factor with a binding site at -881 bp in the human *UCP1* promoter, although not detected at a stringent probability level (p<1e-5).

**Table 2.**
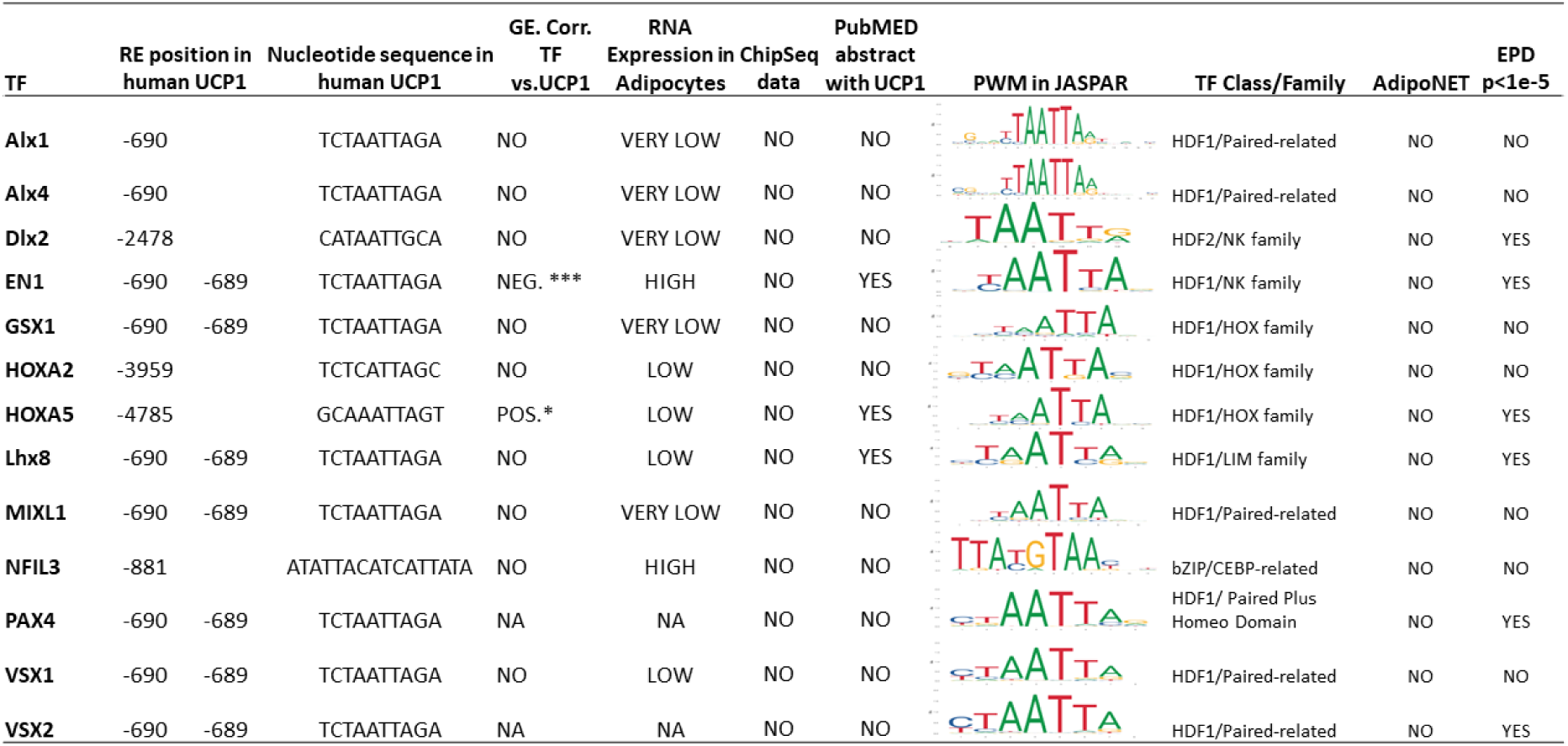
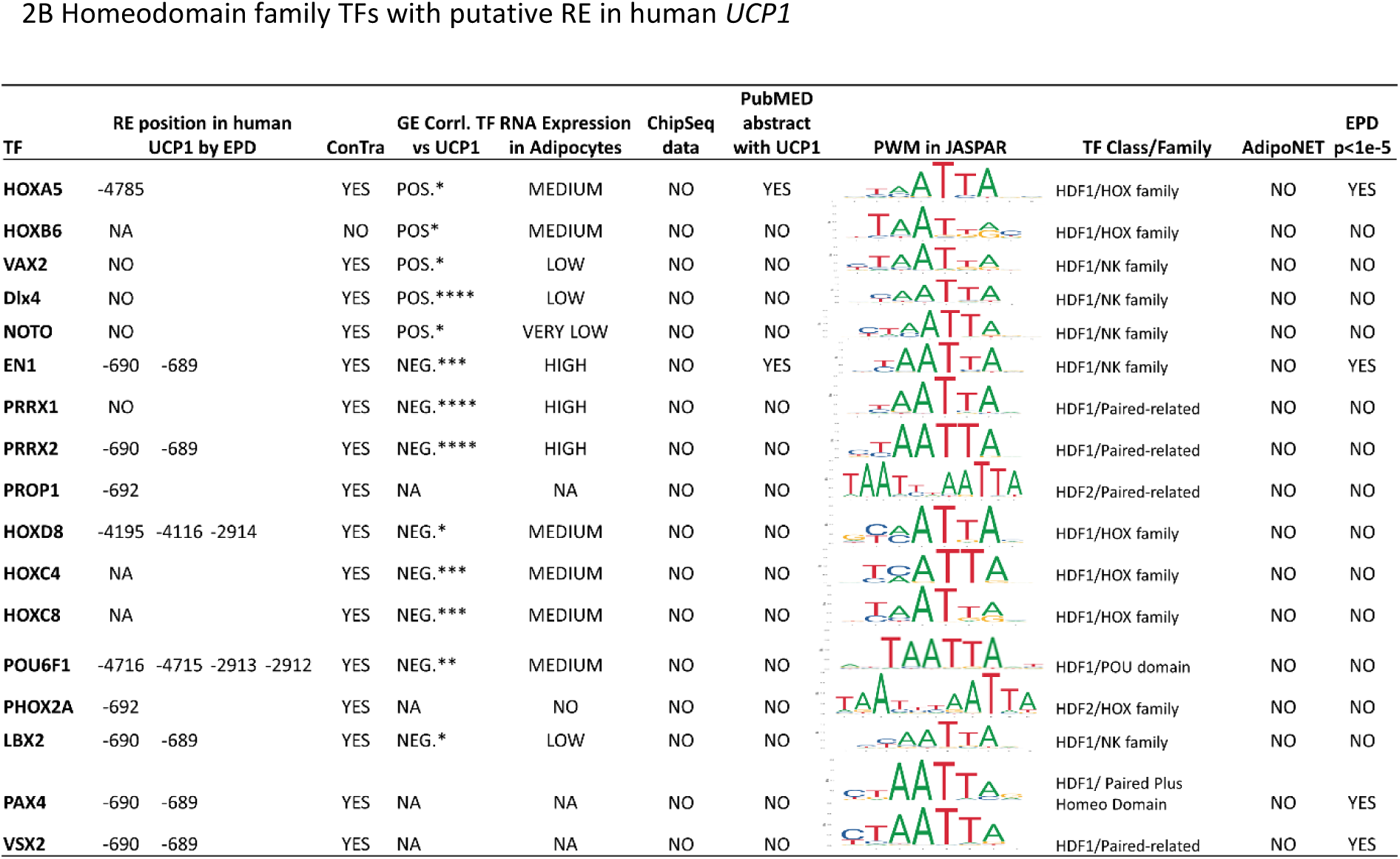
2A Human-specific *UCP1* putative TFs. Tables 2A and 2B summarize our data on TFs with an RE in the human *UCP1* promoter identified by EPD (p<1e-4) and filtered according to specific criteria. These include the position of the RE relative to the TSS, their gene expression pattern relative to *UCP1*, their gene expression levels measured in neck-derived preadipocytes and *in vitro* differentiated adipocytes, their ChipSeq data from the ChiPBase and TFlink databases, their classification into TF class and family based on JASPAR databases, their PWM, their involvement in the adipocyte phenotype formation pathway according to the AdipoNET interface, EPD TF RE search results at p<1e-5 significance level in the human *UCP1* promoter. Bold highlights TF or value outstanding in the related category. Table 2**A** highlights the 13 putative regulatory TFs that possess REs only in the human *UCP1* promoter and are absent from the other *UCP* genes examined. Table 2**B** summarizes our data for 17 TFs belonging to the homeodomain Family 1 and 2 TF classes whose RE matches the human-specific binding sequence identified in the *UCP1* promoter ("TCTAATTAGA") the expression of which is correlated with *UCP1* or for which we do not have gene expression data and therefore cannot exclude them. GE: Gene Expression, Corr.: Correlation.

On the other hand, 93 TF REs were consistently present in all *UCP* promoters of the three species analysed, indicating some degree of shared regulation (Figure 2F Purple box, Suppl Table 1). These include TFs for which phylogenetically well-conserved binding sites were found in the *UCP1* promoters in previous studies, such as PPARA, PPARG, RARE, TFAP2, SP, THR and SREBF1 (Gaudry and Campbell 2017).

#### 2.3 Narrowing transcription factors that bind to the unique palindromic nucleotide sequence of the human UCP1 regulatory region

Next, we wanted to predict the TFs that could bind to the unique nucleotide sequence "5’- TCTAATTAGA-3’" located at -690 bp in the human *UCP1* regulatory region. In addition to the 9 human-specific TFs from the homeodomain family, the Eukaryotic Promoter Database also revealed several TFs from the homeodomain family (p < 1e-4) that potentially recognise this RE but lack specificity for human *UCP1* (Suppl Table 2). Therefore we performed a comprehensive analysis to determine which of the proposed TFs are likely to play a functional role and to assess their importance in the adipocyte browning process (Table 2B). First, we verified the presence of the predicted TF REs of homeodomain family 1 and 2 (HDF1-2) using the ConTra v3 platform (stringency: core=0.95; similarity matrix=0.85; http://bioit2.irc.ugent.be/contra/v3/#/step/1) (Suppl Fig. 3).

In addition to confirming the results of the Eukaryotic Promoter Database, the ConTra v3 interface can be used to gain insight into the evolutionary conservation of REs based on aligned sequences from many species. Contra v3 analysis verified the set of TFs potentially associated with the *UCP1* promoter at -690 bp predicted by the Eucaryota Promoter Database, as the HDF TFs were also identified by ConTra v3 (Suppl. Fig 3.). Therefore, we extended our investigation to all TFs of the HDF1 class based on JASPAR, as they could potentially bind to the *UCP1* promoter at this position. To narrow down the group of potential homeodomain family TFs, we examined their relative gene expression levels in our neck-derived adipocyte samples and focused on those that showed a significant positive or negative correlation with *UCP1* gene expression (Table 2B; Suppl Fig. 4, Suppl Table 3). For some TFs, we did not have gene expression data, so we could not exclude them from the group of TFs that might bind to this specific RE. We collected additional data on this narrowed group of TFs, including ChipSeq data based on ChiPBase and TFlink databases, search results from the Eukaryotic Promoter Database using more stringent p < 1e-5 criteria, and position weighting matrices (PWMs) from the JASPAR database to see the sequences of the REs (Table 2A, 2B). Moreover, we used the AdipoNET platform (Tóth et al. 2021) to categorise these TFs based on their potential involvement in the formation of brown, white or both adipocyte phenotypes as “linker” pathway genes. This categorization provided insights into the possible roles of these proteins in specific adipocyte phenotypes. In addition, we performed an automated search of the PubMed database to identify the co-occurrence of these TFs and *UCP1* in article abstracts to determine what knowledge we have about these TFs concerning *UCP1* (Suppl Table 4, Suppl Fig. 5.). The data collected indicate that Engrailed Homeobox 1 (EN1), Paired Related Homeobox 2 (PRRX2), Homeobox A5 (HOXA5), Paired Box 4 (PAX4) and Visual System Homeobox 2 (VSX2) are the TFs most likely to bind to the given RE sequence. While EN1 and PRRX2 showed high expression in adipocytes and a strong negative correlation with *UCP1*, HOXA5 showed a positive correlation, but a low expression. Although the expression profiles of PAX4 and VSX2 were not investigated in our previous transcriptomic analyses (Tóth et al. (2020), a more stringent search in the Eukaryotic Promoter Database (p > 1e - 5) also identified REs for these TFs at -690 bp, supporting their potential for functional binding at this position. In addition, the ISMARA platform (Balwierz et al. 2014) shows high activity of the PAX4 motif in adipose tissue (Suppl Fig. 6). Of note, the AdipoNET database and publication data mining so far have not previously linked these homeodomain family TFs with pathways belongs to adipocyte phenotype formation or *UCP1* expression, so we present here a new area for further investigation of the role of these TFs in browning processes.

#### 2.4 Investigation of the vicinity of the human UCP1-specific homeodomain family REs identifies potential regulatory hubs

TFs of the homeodomain family are known to be involved in embryonic developmental processes, suggesting a plausible role in the regulation of tissue-specific *UCP1* expression during adipocyte maturation (Gehring 1994; Banerjee-Basu and Baxevanis 2001). Their recognised function involves the modulation of gene expression through the influence of epigenetic markers that affect the accessibility of chromatin and thereby control the binding of TFs to nearby REs. In addition, homo- or heterodimers are often formed as cooperating complexes to improve their DNA binding specificity (Slattery et al. 2011; Cain et al. 2023). Therefore, our further investigations focused on the regulatory elements of the TFs near the identified human-specific genomic positions (in a range of +40 bp and -40 bp) (Table 2A) to determine which TFs can be allowed to operate by these homeodomain family TFs.

First, we investigated TF REs near position -690 of the human *UCP1* promoter. Using a stringent search (p < 1e-4), the Eukaryotic Promoter Database identified the presence of REs for B-cell lymphoma 6 (BCL6) at -702, regulatory factor X5 (RFX5) at -651/650, and REs for basic helix-loop-helix (bHLH2) class TFs around -679 bp (Fig. 2I). Remarkably, BCL6 and RFX5 showed high expression and a strong positive correlation with *UCP1* in our cervical adipocyte samples (Suppl Fig. 7; Suppl Table 3). Of the bHLH2 TFs with putative RE at -679 bp, ID4, SNAI2 and TCF3/4/12 also showed high gene expression levels but a negative correlation with *UCP1*. Furthermore, the primary interaction network of PAX4 described using the ISMARA platform (Balwierz et al. 2014) also includes TFs identified near its RE at -690 bp, such as BCL6, TCF4, ID4, SNAI2 and RFX5, indicating the potential formation of a regulatory complex (enhanceosome) at this position of the human *UCP1* promoter (Suppl Fig. 8).

Next, we investigated the proximity of other identified human *UCP1*-specific putative REs of the homeodomain family, the REs of HOXA5, HOXA2 and DLX2, located at -4785, -3959 and - 2478 bp, respectively (Table 2A). These REs were present in the primate genomes but absent in the mouse and rat *UCP1* promoters (Table 2A, Suppl. Fig. 3 Contra v3). The HOXA5 and DLX2 REs were also identified by the Eukaryotic Promoter Database with a more conservative probability of p<1e-5. All three TFs are expressed at low levels in adipocytes of cervical origin and only *HOXA5* gene expression showed a positive correlation with *UCP1*. Remarkably, we identified the presence of the HIC1 TF RE near the HOXA5 RE (Suppl Fig 9A), which showed high expression levels in pre-adipocyte samples but a significant reduction in mature adipocytes and a negative correlation with *UCP1* expression (Suppl Table 3). A context- dependent regulatory aspect is suggested by the gene expression profile of TEAD RE, which was detected at -2496 bp near the DLX2 RE (Suppl Figure 9B). TEAD1, TEAD3 and TEAD4, which are highly expressed in neck-derived adipocytes, showed a negative correlation with *UCP1* gene expression, while TEAD2 showed a positive correlation. At the same time, NFKB2, which has a putative RE at -3976 bp, and ZNF410 and ZNF283 with RE at -3947 bp and -3948 bp, respectively, possibly form a regulatory complex with HOXA2 (Suppl Fig 9C). While NFKB2 showed a negative correlation trend with *UCP1* gene expression, ZNF410 and ZNF283 showed no obvious correlation (Suppl Table 3). Importantly, these TF REs detected near the homeodomain family TF REs were also identified at a more stringent search threshold (p < 1e- 5), emphasising the likelihood of their involvement in regulatory complexes.

Additionally, our investigation extended to the vicinity of the unique NFIL3 RE identified at - 881 bp within the human *UCP1* promoter (Table 2A). Here, the presence of the REL RE was detected at -902 bp (Suppl. Figure 9D, 9E) and also confirmed the conservation of these RE in the *UCP1* promoter in primates, whereas their absence in the mouse and rat genomes (Suppl. Figure 9F).

#### 2.5 Common RE of the orthologous UCP1 promoters occur at different genomic locations in humans, mice and rats and show no sequence conservation at a particular genomic position

Examination and comparison of the regulatory components in the human, mouse and rat *UCP1* promoters revealed a number of TF REs present in the human *UCP1* promoter, including ARID3B, ATF7, CREB3, MAF::NFE2, NFE2L2, POU6F1, RXRA::VDR, SOX21, TEAD4 and XBP1, which are absent in the human *UCP2* and *UCP3* promoters, but also have REs in the mouse and rat *UCP1* promoters. This group consists mainly of TFs known to be involved in *UCP1* gene expression, including well-characterised TFs such as CREB/ATF (Cao et al 2001; Liu et al 2019), RXR (Alvarez et al 1995), NFE2 and NFE2L2 (Rim and Kozak 2002). A closer look at their gene expression profiles (Fig. 2J) and the prediction of signalling pathways with AdipoNET suggests that their main function is to promote the white adipocyte phenotype, as shown by the strong negative correlation of POU6F1, TEAD4, XBP1 and NFE2L2 gene expression with *UCP1* (Table 3A). Only RXRA gene expression shows a positive correlation with *UCP1* (Suppl Table 3), suggesting a potential enhancer role in browning processes. Consistent with this, AdipoNET predictions suggest that RXRA, CREB3 and XBP1 may contribute to the development of both white and brown adipocyte phenotypes. Interestingly, only POU6F1 from the family of homeodomain TFs was exclusively associated with *UCP1* (it is absent in the promoter of *UCP2* and *UCP3* in mice and rats), indicating its possible role in the specific regulation of the *UCP1* gene and thermogenesis in these species. This observation suggests the presence of evolutionarily conserved TF REs in the *UCP1* promoter in different species. However, a comparative analysis of the regulatory sequences of UCP1 in different species (https://genome.ucsc.edu/) revealed that most of these REs were detectable in the consensus sequences of primates, but not in mice and rats. This may indicate that these REs are not conserved in the consensus sequence of these species, as they occur at different positions in the UCP1 promoter of mice and rats (Suppl. Figure 10A, 10B; for the positions of the TF REs in the different species, see Suppl. Table 2).

**Table 3.**
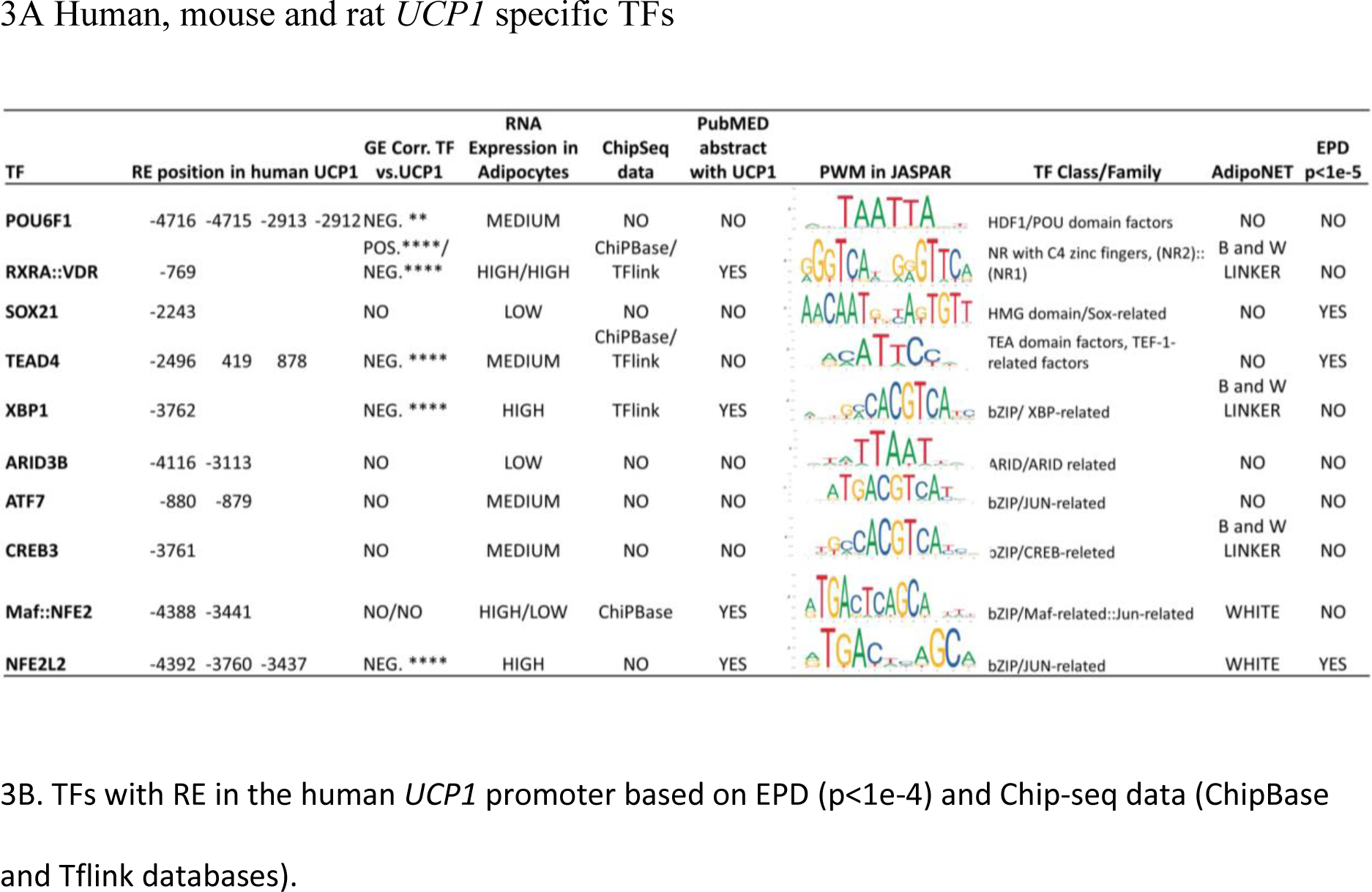
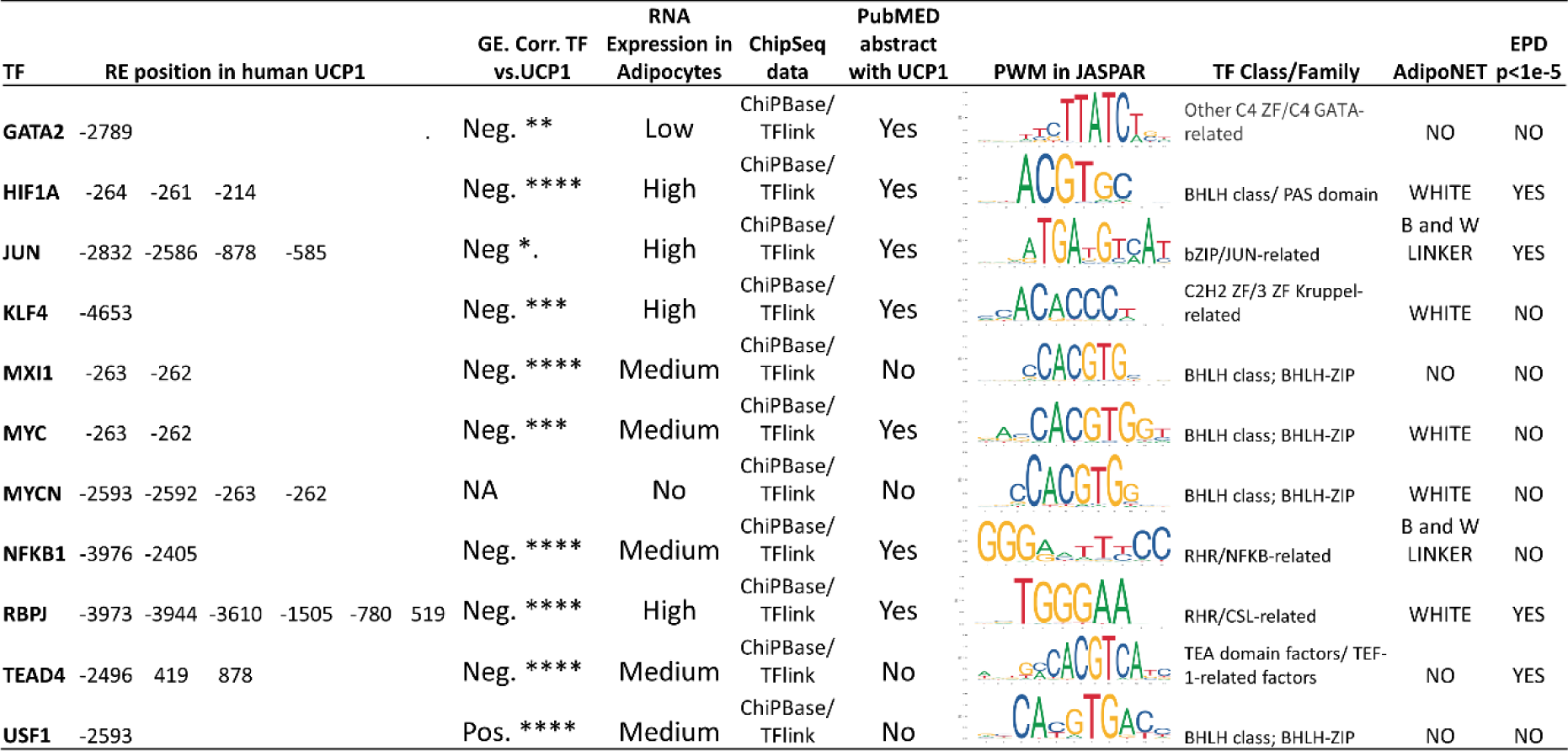
Tables 3A and 3B summarize our data on TFs with RE in the human *UCP1* promoter identified using EPD (p<1e-4) and filtered according to specific criteria. These include the position of their RE relative to the TSS, their gene expression pattern relative to *UCP1*, their gene expression levels measured in neck-derived preadipocytes and in vitro differentiated adipocytes, and their ChipSeq. data from the ChiPBase and TFlink databases, their classification into TF class and family based on JASPAR databases, their PWM, their involvement in the adipocyte phenotype formation pathway according to the AdipoNET interface, EPD TF RE search results at p<1e-5 significance level in the human *UCP1* promoter. Bold highlights TF or value outstanding in the related category. **3A** The table summarizes the 12 TFs with RE found to be common in human, mouse and rat *UCP1.* **3B.** The table summarizes the data of TFs with RE (EPD p<1e-4) in the human *UCP1* promoter that also has ChipSeq data from both ChiPBase and TFlink databases. GE: Gene Expression, Corr.: Correlation.

#### 2.6 Common human-specific bHLH transcription factor response elements in the proximal promoter of UCP1-2-3 genes

Finally, our comparative analysis revealed four regulatory TF elements (HES1, MAX, MNT, TCFL5) that occur exclusively in human *UCP* genes (Suppl Table 1). These TFs have binding sequences within the regulatory domains of the human *UCP1, UCP2* and *UCP3* genes, but are not present in the promoters of the analysed mouse and rat *UCP* genes (Suppl Fig. 11A).

Interestingly, all of these TFs have RE in the proximal promoter region of human *UCP1* at -263 bp. This RE is recognised by the TFs of the basic helix-loop-helix (bHLH) family, and the Eukaryotic Promoter Database assigns additional TFs to this nucleotide sequence beyond the four mentioned above (Suppl Table 2). Therefore, further studies are needed to determine which context-dependent regulatory mechanisms might enable the TFs to actually be users of this RE.

#### 2.7 Predicted and experimentally validated TF binding sites in the human UCP1 promoter show little overlap

**I**n our study, we performed a detailed comparative analysis of putative TF REs identified using the Eukaryotic Promoter Database (https://epd.epfl.ch) in the regulatory region of the human UCP1 gene extending from -5000 to +1000 bases from the TSS, with a significance level of p < 1e-4. We than compared these results with experimentally validated binding proteins/TFs for the same DNA sequence using the ChiPBase (https://rnasysu.com) and TFlink (https://tflink.net/) databases. While we identified 364 putative TF REs using the Eukaryotic Promoter Database, ChiPBase recognised 170 and TFlink 131 experimentally validated TFs (Fig. 3A). Significantly, only a small proportion of the experimentally identified binding TFs had a matching RE recognised by the Eukaryotic Promoter Database (ChiPBase: 23 TFs, 14%; TFlink: 41 TFs, 31%). Conversely, we found that only 53 TFs (15%) of those with REs according to the Eukaryotic Promoter Database were associated with laboratory validation based on these two databases. Note that the two Chip-Seq databases also have only a small overlap. Notably, only 11 TFs were identified in all three databases (GATA2, HIF1A, JUN, KLF4, MXI1, MYC, MYCN, NFKB1, RBPJ, TEAD4, and USF1), highlighting their potential role in regulating human *UCP1* gene expression. The gene expression patterns of these TFs in neck-derived adipocyte samples (Tóth et al. 2020) further support this, with most showing a strong correlation with *UCP1*, mainly negative, while Upstream Transcription Factor 1 (USF1) showed a positive correlation (Fig. 3B). USF1, whose REs were identified in all *UCP*s of the three species studied, appears to act as a general regulator of gene expression of *UCP*s and not as a specific regulator of *UCP1* and browning processes (Suppl Table 2) (Laurila et al. 2016). While the predictions of AdipoNET also suggest that most of these TFs are involved in the development of the white adipocyte phenotype, JUN and NFKB1 may play a role in both white and brown adipocyte phenotypes (Table 3B). Response elements of HIF1A, JUN, RBPJ and TEAD4 were identified at a more stringent significance level (p < 1e-5), supporting their possible involvement in the regulation of *UCP1* gene expression. Interestingly, four of these 11 TFs (HIF1A, MYC, MYCN and MXI1) share a common binding site around -263 bp in the proximal promoter region, which was previously identified in our study as a potential site for fine-tuned, state-dependent regulation of human *UCP* gene expression (Suppl Fig 11A). This RE is not evolutionarily conserved in the mouse and rat *UCP* genes, highlighting the unique control aspects of *UCP1* in humans and the potential for the formation of a regulatory module with TFs SP2 and EGR1, which have REs nearby (Suppl Table 2), based on data revealed by ConTra v3 (Suppl Figs 11B, 11C, 11E).

**Figure 3.**
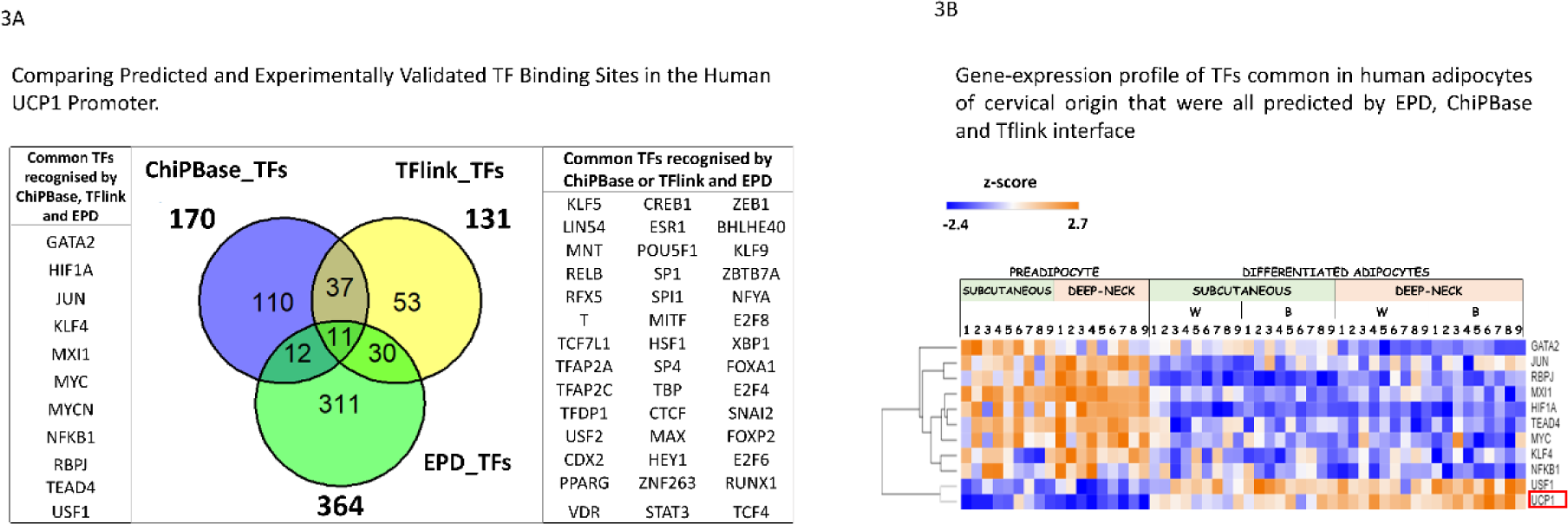
**3A** The Venn diagrams show the number of TFs with putative RE in the human *UCP1* promoter (-5000 – +1000 BP) based on EPD, ChiPBase and TFlink databases. TFs commonly found in the databases are named in the side panels (Venn program in R environment). **3B** The heatmap shows the gene expression profiles of 11 TFs in human adipocytes of cervical origin that have been predicted or experimentally shown to have RE in the human *UCP1* promoter based on EPD, ChiPBase and TFlink interfaces. For comparison, *UCP1* is also shown. Hierarchical cluster analysis based on the Spearman Rank similarity matrix identifies TFs with similar gene expression profiles to *UCP1* (Morpheus interface).

## DISCUSSION

### 1. Evolutionary dynamics of *UCP1* genes and their promoters differ in humans and rodents

#### 1.1 Research trends in the phylogenetic analysis of genes

*In silico* studies based on DNA sequences developed into a fundamental approach for understanding the evolutionary dynamics of gene-specific functions. In general, phylogenetic analyses have primarily focused on coding sequences, and there are now more than a hundred species in comparative analyses of orthologous genes for *UCP1*, providing insights into complex phylogenetic relationships (Cassard et al. 1990; Klingenspor et al. 2008; McGaugh and Schwartz 2017, Hughes et al. 2009; Jastroch et al. 2005, 2008, 2018; Berg et al. 2006; Hou et al. 2017; Gaudry et al. 2017). In the meantime, evolutionary studies on transcriptional regulation have remained strikingly limited. In particular, Gaudry and Campbell (2017) performed a comparative analysis of the transcriptional regulatory elements of *UCP1* in mammals, although they focused only on the REs already described, particularly in the core promoter and distal enhancer. Their work has shed light on evolutionarily conserved key sites such as the TATA box, RARE-3 and PPRE motifs which are expected to be functionally relevant in the phylogeny of eutherians (Gaurdy and Campbell, 2017; McGaugh and Schwartz, 2017). Our studies demonstrate the ancient origin of these regulatory elements, indicating a remarkable level of conservation. According to searches in the Eukaryotic Promoter Database, these elements are widespread in the promoter regions of the *UCP* genes in all three species studied. This abundance may indicates a fundamental role in the regulation of gene expression within this gene family. However, due to their widespread distribution in the *UCP* family, these elements may be less suitable as specific targets for precise modification of the transcription of the *UCP1* gene, which is the ultimate goal of metabolic disorder control. Nonetheless, the results of Gaudry and Campbell (2017) suggest that transcriptional control of gene expression is only conserved to a limited extent within the mammalian clade. This observation suggests the possible existence of novel regulatory and functional elements within the human *UCP1* promoters that may adapt and coordinate to species-specific conditions and enable the development of more targeted interventions.

#### 1.2 Differences between the regulatory region of UCP1 in humans and rodents

Currently, several interfaces are developed for the prediction of DNA binding of TFs with known PWM based on the promoter nucleotide sequence, such as the Eukaryotic Promoter Database we used, TRANSFAC2.0 (https://genexplain.com/transfac-2-0/), GTRD (Yevshin et al. 2017) or ConTra v3. Using these interfaces, the set of potential regulatory sequences identified in the 9 UCP promoters and their putative associated TFs supports the possibility of distinct regulation of gene expression in these species, as similarity analysis of RE profiles reveals differential clustering between human and rodent UCPs. In particular, the regulatory profile of *UCP1* in humans shows the closest proximity to *UCP2* in humans, whereas *UCP1* in mice shows a closer relationship to *UCP1* in rats. Our comparative analysis of upstream regulatory segments in human, mice and rats *UCP1* genes reveals the intricate interplay between sequence conservation and divergence that characterises the evolutionary trajectory of these genes. While a substantial proportion of the identified cis-acting RE in these segments of rodent and human *UCP1* genes are common (174 of 364; Suppl Figure 2B), broadly consistent with previous findings (Collins et al., 2004), our analysis of sequence similarity reveals major differences (Figure 1B), suggesting the possibility of the emergence of distinct regulatory modules (Saito et al. 2008). Remarkably, a significant portion of the distal enhancer region of the human *UCP1* promoter is missing in the mouse and rat genomes. This may be due to secondary losses in the ancestors of mice and rats in the DNA segment that arose after the opossum lineage split. Remarkably, these DNA sequence losses did not lead to extinction of the rodent ancestors, suggesting that their existence was not essential for life or that possible compensatory mechanisms allow sufficient TF binding to other parts of the regulatory region. Despite this sequence deletion, human-specific *UCP1*-REs were identified in the DNA regions whose consensus sequences are found in rodents, which could potentially form the basis for human-specific UCP1 regulation. Recent studies also suggest interspecies variability in the regulation of *UCP1* expression. They show that the 3’-untranslated region (3’-UTR) of *UCP1* mRNA is processed differently in mice and humans, which quantitatively affects *UCP1* synthesis and thermogenic activity (Lu et al., 2021). These differences urge caution when interpreting research results obtained in rodent models to humans.

#### 1.3 Repeat elements in the human UCP1 promoter

We identified numerous repeat elements in the human *UCP1* promoter region, including those embedded in evolutionarily novel sequences that are absent in the corresponding rodent regulatory regions (Fig. 1B). BLAST searches revealed the presence of these repeat sequences at several locations in the human genome (Fig. 1H). However, the mechanisms underlying their integration into a genome remain unclear, in particular whether these mobile elements are randomly integrated or controlled by specific sequences. We observed the presence of short palindromic sequences at the ends of the novel DNA segments, some of which are annotated as TF REs in the Eukaryotic Promoter Database. Palindromic sequences in DNA are traditionally recognised as specific binding sites for nucleotides or proteins, leading to functional consequences (Pingoud and Jeltsch 2001; Linheiro and Bergman 2008). This may suggest that these binding motifs could be ancient genetic elements used for integration into the genome and repurposed by eukaryotic cells as regulatory modules to control gene expression (Thompson et al. 2016). For example, the multiple integration of the MLT1G repeat sequence into the human genome may organise the expression of nearby genes into a regulatory circuit, leading to the simultaneous expression of the genes involved. This could facilitate the development of complex processes such as adipocyte thermogenesis.

An interesting observation related to the 5’-TCTAATTAGA-’3 palindromic homeodomain RE sequence identified at -690 bp in the human *UCP1* promoter may also support the repurposing hypothesis. The same nucleotide sequence was previously recognised by Li and co-workers (2005) as a characteristic DNA fragment located at one endpoint of the 1165 bp long sequence that distinguishes two genotypic variants of Mamestra configurata nucleopolyhedrovirus-A (MacoNPV-A). These variants include the lower virulence variant 90/4 and the archetypal variant 90/2. They concluded that the recognised palindrome sequence raises the possibility that the difference between these viral variants is due to the insertion or deletion of a transposable element. RepeatMasker does not recognise this particular sequence region as a repetitive element in humans, but mutational changes since the ancient insertion may have masked the repetitive element.

Other studies also suggest that transposable elements and short tandem repeats may influence the regulation of gene expression in the genome (Horton et al., 2023; Chuong et al., 2016; Kunarso et al., 2010). For example, Scarpato and colleagues (2015) have suggested that some repetitive elements, such as SINEs, are highly prone to hybridisation with microRNAs, thereby modifying the transcriptional activity of nearby genes. Overall, our results suggest that species-specific differences in transposable and repetitive elements in the *UCP1* promoters may underlie gene-specific regulatory mechanisms.

### 2. Comparative analyses of putative RE in *UCP1, UCP2* and *UCP3* promoters in humans, mice and rats underline the possible regulatory role of TFs of the homeodomain family

#### 2.1 The human UCP1-specific 5’-TCTAATTAGA-3’ binding sequence

One of the main goals of our study was to identify unique REs within the human *UCP1* promoter which may have remained hidden in "exploratory" studies commonly performed in rodents. Results indicate that most of the known putative regulatory factors of human *UCP1* are general regulators of the studied *UCP*s, which classifies them as less specific targets. Uncovering the specific regulatory elements allows for targeting metabolic processes by safely promoting fat cell browning. Our study focused on the identified unique TF REs of the homeodomain family of human *UCP1* and their vicinity, in particular a palindromic nucleotide sequence, 5’-TCTAATTAGA-3’, located at -690/689 bp in the human *UCP1* promoter (Fig. 2I, Fig. 4). It confers the ability of homeodomain family TFs to bind to both DNA strands, allowing dimerisation and stronger tethering (Jolma et al. 2013; Cain et al. 2023; Dror et al. 2014). It is also possible that multiple TFs of the homeodomain family alternate the use of this binding site depending on the circumstances, and these TFs could differ in whether they promote or block gene expression (Foulkes et al. 1991). The homeodomain family is one of the largest groups of TFs in metazoans and consists of almost 200 family members in humans (Lambert et al. 2018). Their recognised functions include the control of developmental and homeostatic processes through the modulation of DNA availability (Waldén et al. 2012; Hilton et al. 2015; Cantile et al. 2003), but their potential for species-specific regulation of adipocyte phenotype remodelling has been underestimated. These TFs can bind very similar AT-rich monomer sequences, with small differences in nucleotides being sufficient for a specific interaction and modulating the openness of DNA to allow or deny TFs access to nearby REs (Park et al. 2009). In our analysis, regulatory elements such as the bHLH type 2 TFs, BCL6 and RFX5, identified near this palindromic sequence unique to human *UCP1*, may further increase the likelihood of binding by forming a complex/enhanceosome with homeodomain family TFs via a process termed "epistatic capture” (Emera and Wagner 2012). The combined effect of these factors can lead to a regulatory output that is not simply additive but is influenced by the specific combination of transcription factor binding. Furthermore, in a previous study analysing the signalling pathways enriched by genes with differential expression in subcutaneous and deep neck samples, both in preadipocytes and differentiated mature adipocytes, the homeodomain family TF pathway was found to significantly dominate in subcutaneous adipocytes, indicating their possible active involvement in the regulation of browning processes (Tóth et al. 2020).

**Figure 4.**
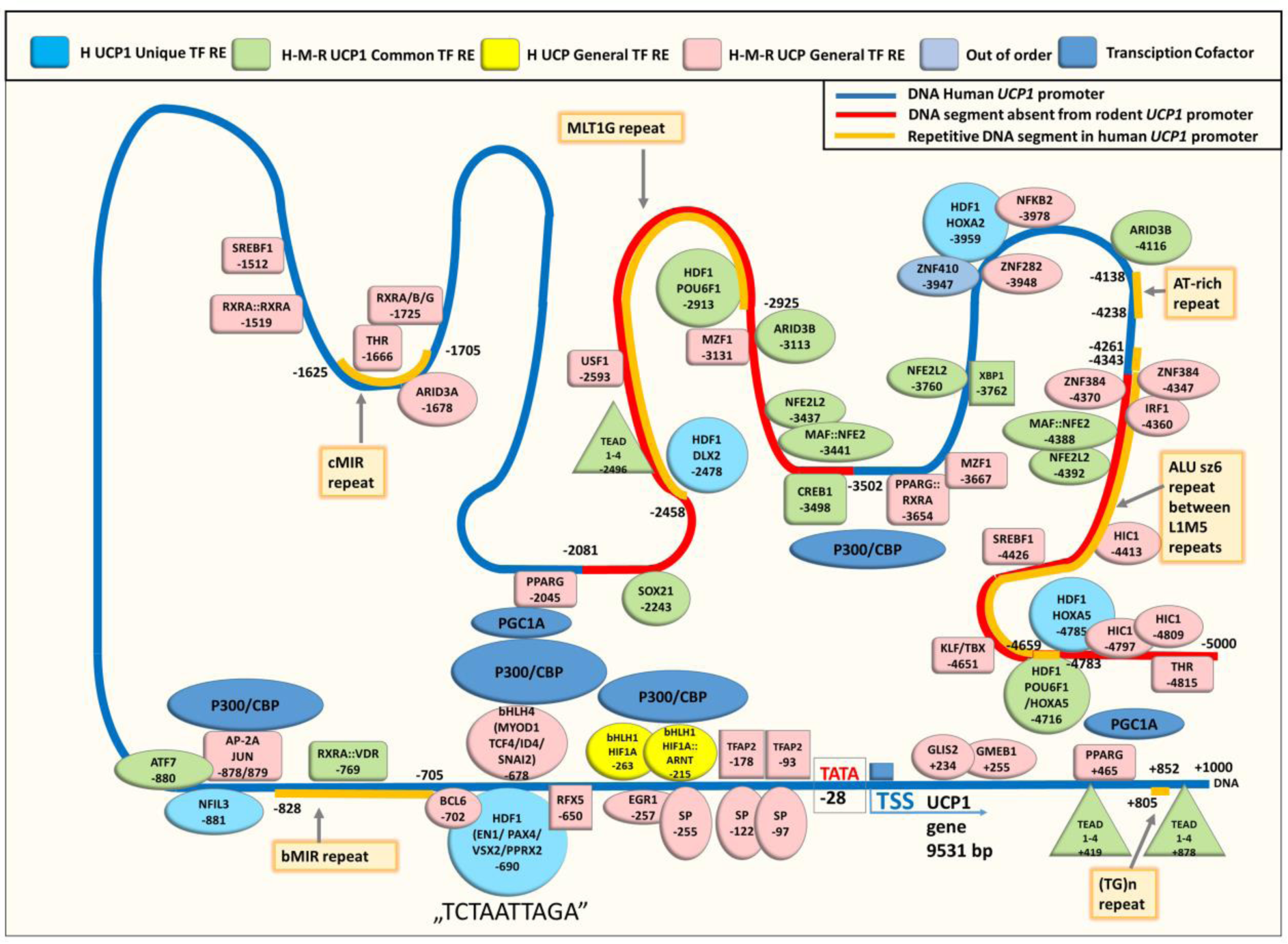
This figure provides a comprehensive overview of the transcription factor response elements identified and filtered by our study within the human *UCP1* promoter. These REs exhibit gene or species specificity. Additionally, several previously reported TF REs, which did not display such specificity, are included for reference. The figure highlights specific promoter segments in the human *UCP1* promoter that are absent in the mouse and rat *UCP1* promoters (indicated by the red line). The yellow line represents repeat elements identified by RepeatMasker. Numerical annotations within the promoter denote base pair positions relative to the transcription start site (TSS). For clarity, a list of TF abbreviations is provided in the "List of abbreviations" section at the end of the article. H: human, M: mouse, R: rat. HDF: homeodomain family, bHLH: basic helix-loop-helix family (PowerPoint).

#### 2.2 Prediction of transcription factors targeting the human-specific sequence ‘5’- TCTAATTAGA-3’"

Among the TFs of the homeodomain family, we propose several potential binding candidates for the human UCP1-specific RE at -690 bp. EN1, which is abundant in human neck adipocytes and whose gene expression negatively correlates with *UCP1*, has previously been shown to be a positive regulator of brown adipogenesis in mice (Zhang et al. 2016; Singh et al. 2015; Atit et al. 2006), but was downregulated after animals were exposed to cold. These studies show that EN1 is a marker for the brown adipocyte lineage, suggesting a role in brown adipocyte development, but not necessarily a direct regulator of *UCP1* expression during thermogenic activation in rodents. Our previous human study has shown that it is upregulated in the obesity phenotype in adipocytes carrying the single nucleotide polymorphism (SNP) FTO rs1421085 T-to-C (Tóth et al. 2020, unpublished data), suggesting a diverse, species-specific role in the regulation of *UCP1* expression. In contrast, another candidate, HOXA5 has been reported to be increasingly expressed in the thermogenic fat cells of rodents and to play a role in promoting adipose tissue browning through anti-inflammatory processes (Cao et al. 2018; Kato et al. 2021; Singh et al. 2016). Consistent with this study, HOXA5, showed a gene expression profile which may suggest its beneficial effect on browning in cervical-derived human adipocytes (Fig. 2H, Suppl Table 3)., At the same time, other human study showed increased expression in white adipocytes derived from induced pluripotent cells (Mohsen-Kanson et al. 2014). This discrepancy also suggests that HOXA5 may play a context-dependent role in the expression of the adipocyte phenotype in different species and fat depots.

The known role of PAX4 in pancreatic beta cell development and function is associated with the maintenance of glucose homeostasis (Brun et al. 2008; Xu et al. 2017), and abnormalities in its expression have been linked to metabolic diseases such as diabetes (Lau et al. 2023). However, its role in adipocyte function is less understood. PAX4 is expressed in mesenchymal progenitor cells of adipocytes (Xu et al. 2017), and our study suggests that it may be involved in the regulation of adipogenesis, including the balance between white and brown adipocyte formation (Suppl Fig. 6, Fig. 8). Role of the other candidate TFs of the homeodomain family, PRRX2 and VSX1/2, in the context of the thermogenic response has not yet been characterised. Nonetheless, single nuclear RNA sequencing studies have identified PRRX2 with differential expression activity in brown versus white adipogenesis in human cells (Gupta et al. 2023), indicating a possible role in thermogenic adipocyte formation.

The TF REs identified in the human *UCP1* promoter are often evolutionarily conserved in primates, but not in mice and rats. Even the identified common TF REs in orthologous *UCP1* promoters, such as POU6F1 TF from the homeodomain family, are located in the promoter segment in humans that is absent in mice and rats and are not conserved in the genomic consensus sequences. This indicates a kind of "mobility" of the regulatory elements and possible convergent evolutionary processes. Nonetheless, according to our data mining efforts, POU6F1 has not been previously associated with adipocyte function. Recent research has identified its role in lung biology and cancer-related processes, revealing its interaction with RORA in the lung leading to downregulation of *HIF1A* gene expression and potential inhibition of adenocarcinoma cell proliferation (Xiao et al., 2022). However, as shown in previous studies (Takahashi et al., 2007), certain POU factors serve as effective mediators of cell reprogramming. Given its strong expression in mature adipocytes and a remarkable negative correlation with *UCP1* gene expression (Table 3A, Suppl Table 3), it is plausible that POU6F1 may play a role in adipocyte phenotype switching and that this function is conserved in mammalian evolution.

#### 2.3 Limitations in the identification of functional regulatory elements in the human UCP1 promoter

Our comparative analysis revealed little overlap between the TF REs predicted in our study based on DNA sequence analyses and those identified experimentally with ChipSeq in the human *UCP1* promoter, highlighting the challenges in identifying functional regulatory elements. TFs, like other proteins, have the capacity for non-specific DNA binding, which is essential for efficient target site localisation by facilitating linear diffusion (Pingoud and Jeltsch 2001). This non-specific binding usually involves interactions with the DNA backbone and not with the bases. In contrast, specific binding is characterised by a close interplay between direct (interaction with the bases) and indirect (interaction with the backbone) selection. The frequent occurrence of non-functional TF binding can lead to considerable ambiguity about functional DNA-TF binding in ChipSeq laboratory experiments. In addition, the infinite combinations of biological tissues, developmental stages and specific conditions significantly affect *ad hoc* TF binding to specific DNA segments, making generalisation of results difficult. While PWMs are useful for predicting functional TF binding within promoter sequences, they also have numerous limitations. One of these is the random occurrence of short binding sequences in the genome, which emphasises the need for rigorous statistical analysis to distinguish between biologically meaningful TF binding events and random matches. In addition, the use of species- and gene-specific screening approaches can further increase the reliability of predictions. The used Eukaryotic Promoter Database employs robust statistical tests to predict PWM in a DNA sequence and thus provides the best possible prediction. Other limitations of the *in silico* approach are our incomplete knowledge of the binding sequences of TFs and the inability of PWMs to fully capture the sequence flexibility of TF binding sites. TFs can tolerate variations in DNA sequence at their binding sites, leading to a degree of "sloppiness" or degeneracy. This sequence flexibility allows TFs to recognise and bind a range of related sequences which may differ from the consensus motif defined by the PWM. Therefore, PWMs may miss functional TF binding sites that differ from the consensus sequence or erroneously predict non-functional sites that are very similar to it. In addition, the PWMs do not take into account the context-dependent nature of TF binding, where neighbouring DNA sequences or chromatin structures may influence binding specificity. As a result, both predictions may miss biologically relevant binding events or lead to false positives. This highlights the importance of integrating experimental and complementary computational approaches to improve the accuracy of TF binding site identification.

### 3. Hotspots in human UCP1 promoter

The prediction of human *UCP1*-specific TFs identified using the Eukaryotic Promoter Database indicates specific DNA regions with higher TF-RE density within the human *UCP1* promoter (Fig. 4). Such an accumulation of regulatory binding sites may lead to the formation of enhanceosomes, in which molecular elements such as TFs and cofactors form complex multilevel regulatory networks that finely orchestrate gene expression. In particular, in the enhancer region of the human *UCP1* promoter, novel regulatory cluster-forming nucleotide sequences are observed at various homeodomain family binding sites, often located at the beginning or end of DNA regions identified as repetitive elements. In addition, several REs appear in the proximal promoter near the RE of the bHLH TF family identified at -262/-263 bp, suggesting the possibility of regulatory complex formation (see Suppl. Figs 11B, 11C, 11D). The proximity of EGR1 and SP2 binding sites provides a scenario for dynamic regulation that could occur through interaction with HIF1A, modulating *UCP1* expression according to cellular conditions. This proposed mechanism shares similarities with mechanisms described in non-small cell lung cancer, where interactions between HIF1A and EGR1 as well as HIF1A and SP1 play a central role in regulating *EPOR* gene expression in response to changing hypoxic conditions (Su et al. 2019). Given the known role of hypoxia and HIF1A in adipose tissue function, the possible involvement of HIF1A in *UCP1* transcription through promoter binding and interaction with other TFs could be an interesting area of research (Tóth et al. 2021; Basse et al. 2017). In addition, EGR1 has already been shown to be involved in metabolic diseases and can influence the insulin sensitivity of fat cells and decide on PI3K/Akt and MAPK signalling pathways (Zhang et al. 2013). It is conceivable that slight variations in the concentrations of different components within the *UCP1* enhanceosomes may interact synergistically and lead to significant differences in *UCP1* expression, as has previously been hypothesised in comparative studies in mice (Xue et al. 2005). Among the potentially related cofactors, the p300/CBP-associated factor with well-described histone acetyltransferase activity stands out as a frequently described partner of TFs with a putative RE in the human *UCP1* promoter, indicating an essential role for them in shaping the fat cell phenotype (Fig. 4). Nevertheless, PGC1A, another described coactivator of TFs that regulate *UCP1*, such as PPARG, appears to interact with TFs that lack *UCP* gene or species specificity (Figure 4). Determining the key components of the regulatory complexes, describing their role and identifying the components that can be replaced or omitted in the initiation of human UCP1 transcription under certain conditions is a promising future research direction.

## METHODOLOGY

### 1. Reconstruction of the Phylogenetic Trees

To generate a combined human, mouse and rat UCP1, UCP2, and UCP3 amino acid, coding sequence and promoter sequence phylogenetic tree, the amino acid and the mRNA data set was downloaded from the National Library of Medicine (https://www.ncbi.nlm.nih.gov), whereas the sequences of gene promoters were downloaded from the EPD database (link: https://epd.epfl.ch/). Accession numbers of the promoters’ FASTA sequences: human *UCP1* FP007435, *UCP2_1* FP016503, *UCP2_2* FP016504, *UCP3_1* FP016505, *UCP3_2* FP016506; mouse *UCP1* FP011827, *UCP2* FP010492, *UCP3* FP010490; rat *UCP1* FP011518, *UCP2* FP000767, *UCP3* FP000766; Accession numbers of the mRNAs’ FASTA sequences: human UCP1 NM_021833.5, UCP2 NM_001381943.1, UCP3 NM_003356.4; mouse UCP1 NM_009463.3, UCP2 NM_011671.6, UCP3 NM_009464.3; rat UCP1 NM_012682.2, UCP2 NM_019354.3, UCP3 NM_013167.2; Accession numbers of the amino-acids’ FASTA sequences: human UCP1 NP_068605, UCP2 AAC51336, UCP3 AAC51367.1; mouse UCP1 NP_033489, UCP2 AAB17666, UCP3 AAB87084; rat UCP1 NP_036814, UCP2 BAA25698, UCP3 AAB71523. The resulting amino acid, exon and promoter sequences were aligned using MAFFT 7.505 (Katoh & Standley, 2013), and a maximum likelihood tree was constructed using IQtree2 2.0.7 (Minh et al., 2020) with automatic model selection (‘-m MFP’) turned on and using 1 000 aLRT and 1 000 rapid bootstrap replications to test the robustness of the analysis. The alignment of the mRNA sequences had a length of 3,398 bp, of which 1536 were variable and 878 informative, whereas the promoters had 7,956 variables and 5,412 informative sites on an alignment length of 12585 bp. Of the 313 aligned amino acid positions, we identified 172 polymorphic and 150 informative characters. Phylogenetic trees were visualized with packages ape 5.7.1 (Paradis & Schliep, 2019) and phangorn 2.11.1 (Schliep, 2011) using R 4.2.2 (R Foundation for Statistical Computing, n.d.). Pairwise genetic distances of *UCP* promoters and exons were calculated with the R package pegas 1.2 (Paradis, 2010) using the “K80” model of nucleotide substitution (Kimura’s 2-parameters distance metric; Kimura, 1980). Pairwise distances were visualized as a neighbour-joining phylogram with pegas, as a UPGMA dendrogram with phangorn, and as a heatmap with pheatmap 1.0.12 (https://cran.r-project.org/web/packages/pheatmap/index.html).

### 2. 3D Protein Structure Prediction, Alignment and Pairwise Structural Comparison

The 3D structures of human, mouse and rat UCP1, UCP2 and UCP3 proteins were downloaded using the AlphaFold 2 protein structure prediction interface (https://alphafold.ebi.ac.uk/) developed by Deep Mind (Jumper et al. 2021). Colors indicate model confidence, AlphaFold produces a per-residue confidence score (pLDDT) between 0 and 100. Some regions below 50 pLDDT may be unstructured in isolation. Dark blue: Very high (pLDDT > 90); Light blue: Confident (90 > pLDDT > 70); Yellow: Low (70 > pLDDT > 50); Red: Very low (pLDDT < 50).

For alignment of the predicted protein structures we used the RCSB PBD (https://www.rcsb.org; Berman et al. 2000) 3D protein structure comparison interface. For protein comparisons, pairwise comparisons were performed with human UCP1. The metrics used to describe the degree of pairwise structure alignment and structural similarity are as follows:

**RMSD** (root mean square deviation) is computed between aligned pairs of the backbone C-alpha atoms in superposed structures in Å resolution. The lower the RMSD, the better the structure alignment between the pair of structures. This is the most commonly reported metric when comparing two structures, but it is sensitive to local structure deviation. If a few residues in a loop are not aligned, the RMSD value is large, even though the rest of the structure is well aligned.

**TM-score** (template modeling score) is a measure of topological similarity between the template and model structures. The TM-score ranges between 0 and 1, where 1 indicates a perfect match and 0 is no match between the two structures. Scores < 0.2 usually indicate that the proteins are unrelated while those > 0.5 generally have the same protein fold (e.g., classified by SCOP/CATH).

**Identity** (sequence identity percentage) is the percentage of paired residues in the alignment that are identical in sequence.

**Equivalent Residues** is the number of residue pairs that are structurally equivalent in the alignment.

**Sequence Length** is the total number of polymeric residues in the deposited sequence for a given chain.

**Modeled Residues** is the number of residues with coordinates that were used for structure alignment.

### 3. BLAST Search to Identify Specific Nucleotide Sequences in the Human, Mouse and Rat Genome

The BLAST interface (https://blast.ncbi.nlm.nih.gov/) was used to identify sequences similar to a given nucleotide sequence in the human, mouse and rat genomes (Altschul et al. 1997). The human RefSeqGene, rat mRatBN7.2 and mouse GRCm39 reference genome sequences were used in the searches. The graphical summary of the results in Suppl. Fig. 1H shows the percentage of homology in different colors, while the position indicates the sequence region of the analysed DNA segment in which the homologous segment is located, showing the hits in separate rows.

### 4. Sequence Presence Assessment for Estimating Evolutionary Divergence

We used the phylogenetic tree based on the genome of the species provided by the Multiz Alignment of 100 Species interface integrated into the UCSC genome browser. We then manually checked the presence of the 1419 bp sequence missing in the mouse and rat genomes in the corresponding sequences of the other species. In this way, we estimated the evolutionary divergence based on the available genomic data and the presence of the sequence in the different species.

### 5. Computational Identification of Putative Transcription Factor Response Elements in Gene Regulatory Sequences Using the EPD Interface

The identification of putative TF REs within the regulatory sequences of the genes under study was performed using the Eukaryotic Promoter Database (EPD) interface (https://epd.epfl.ch) (Périer et al. 2000). EPD’s approach to identify TF REs relies on the JASPAR Transcription Motifs Library (specifically JASPAR CORE 2018 vertebrates) (Sandelin et al. 2004), which includes a diverse set of TF motifs, including generic promoter motifs such as the TATA box, Initiator, GC box and CCAAT box. This library contains a large body of information represented as a position weight matrix (PWM) for 579 TF-binding sequences. Many approaches have been developed to model and learn a protein’s DNA-binding specificity, however, comparative analyses indicate that simple models based on mononucleotide PWMs trained by the best methods perform similarly to more complex models (Weirauch et al. 2013).

To determine whether a given binding sequence is present within the regulatory sequence of the target gene, the EPD interface uses a probability-based algorithm with four different p- value thresholds (p-value < 0.01, 0.001, 0.0001 and 0.00001). In this study, we focused on the two most stringent p-value thresholds (p < 0.0001 and p < 0.00001), as empirical evidence from the literature suggests that more permissive probabilities can lead to high false positive rates (Benita et al., 2009). However, our exploratory studies showed that for some genes with experimentally proven regulatory TFs, the software accurately identified the binding sequence only at p < 0.0001, as exemplified by the occurrence of the HIF1A TF RE for genes such as *VEGFA* and *GLUT1*. In addition, considering that every 4000 nucleotides, a given RE of average length 12 bp can be expected to be found randomly (Wunderlich and Mirny, 2009), the identified occurrence itself has limited meaning. It rather suggests that TFs for which we did not identify the RE are less likely to be involved in the regulation of *UCP* gene expression. Consequently, we presented our findings primarily utilizing the less stringent p < 1e-4 criteria. To mitigate the problem of false positives, we applied additional screening methods to filter out less relevant REs and emphasise potential species- and gene-specific TF-REs. Although a stricter threshold (p < 1e-5) would reduce the number of false positives, it would also increase the number of false negatives, potentially omitting significant findings. False-positive results can be managed through further filtering and by collecting more detailed information. In choosing the significance level (p < 1e-4), we were guided by the possibility of comparing the results of similar tests from paralogous and orthologous UCP promoters and thus filtering out false positives in the analysis. However, for reasons of comparison and transparency, we also present results with a more stringent threshold (p < 1e-5) in the tables.

In this study, we analysed regulatory sequences 5 kb upstream and 1 kb downstream of the TSS for the genes under study (eg.: human *UCP1*: GRCh37/hg19; Chr. 4: 141,489,115- 141,495,114; TSS: Chr. 4:141,489,115). The relevant base-pair boundaries of the upstream regulatory DNA segment may vary among genes, therefore the widest DNA region allowed by the EPD interface was analysed. Data retrieval was automated using a custom script, which we have made publicly available on GitHub, increasing transparency and reproducibility.

### 6. Comparative Analysis of the Putative TFs in the Regulatory Sequences of Human, Mouse and Rat *UCP1*, *UCP2* and *UCP3* Genes

Data analysis and screening were conducted in an R interactive environment. Initially, data were extracted from the EPD (https://epd.epfl.ch) interface as individual data files (text files), corresponding to each target gene. EPD utilizes the JASPAR database, which contains PWMs of TF REs for examining their presence within specific DNA sequences (https://jaspar.genereg.net). The EPD database contains two promoter sequences for human UCP2 and UCP3, as these genes have alternative transcription start sites, with the TSSs of the two variants differing by a few 100 bp. Since we did not want to favour either variant, both variants were included in the analysis. Subsequently, the data extracted for each gene were consolidated into a unified database that compiled occurrence information for the 579 TF binding sequences within the promoter regions of three genes: *UCP1*, *UCP2*, and *UCP3* in humans, mice, and rats. This resulting matrix was then utilized to identify distinct clusters of TF REs based on specific criteria. These criteria included identifying TF REs that were specific to human *UCP1* (not present in human *UCP2* or *UCP3* but potentially present in mouse and rat *UCP1*, *UCP2*, or *UCP3*), unique to human *UCP1* (exclusive to human *UCP1*, absent in human *UCP2*, *UCP3* as well as in mouse and rat *UCP1*, *UCP2*, and *UCP3*), specific to *UCP1* (present in both human and mouse and rat *UCP1*, absent in human *UCP2* and *UCP3* but potentially present in mouse and rat *UCP2* and *UCP3*), common to human, mouse, and rat *UCP1* (absent in human, mouse, and rat *UCP2* and *UCP3*), and general human *UCP* TF REs (occurring in both human *UCP1* and *UCP2* and *UCP3* but not in the promoter regions of mouse and rat *UCP1*, *UCP2*, and *UCP3* genes).

### 7. Chromosome position of the investigated *UCP1*, *UCP2* and *UCP3* promoters and genes in the human genom

*UCP1* promoter in human: GRCh 37/hg19 Chr 4:141,489,115 – 141,495,114 size: 6000 bases; *UCP1* gene in human: GRCh 37/hg19 Chr 4:141,480,585 – 141,490,115 size: 9531 bases, minus strand; *UCP2* promoter in human: GRCh 37/hg19 Chr11:73,693,248 - 73,699,247 size: 6000 bases; *UCP2* gene in human: GRCh 37/hg19 Chr11:73,685,717 - 73,694,247 size: 8531 bases; *UCP3* promoter in human: GRCh 37/hg19 Chr11:73,719,130 - 73,725,129 size: 6000 bases; *UCP3* gene in human: GRCh 37/hg19 chr11:73,711,322 - 73,720,130; size: 8809 bases.

### 8. Mapping The Regulatory DNA Regions of the Identified Putative TF REs in Human UCP1 and Comparing them with Aligned DNA Sequences of Other Species

During the course of the study, a group of TF REs with a distance between the filtered out TF REs below +/- 40 bp were identified, potentially indicating a complex regulatory mechanism. This clustering of TF binding sites aligns with a common phenomenon observed in numerous regulatory regions of multicellular eukaryotes, as described in previous studies (Wunderlich and Mirny 2009). Using the Contra v3 interface (http://bioit2.irc.ugent.be/contra/v3/; Kreft et al. 2017) and the UCSC browser (http://genome.ucsc.edu/), putative human *UCP1* TF REs identified by EPD and filtered by comparative analysis of orthologs and paralogs were examined for nucleotide sequence at the defined positions. The results are fully linked to NCBI (http://www.ncbi.nlm.nih.gov/) and UCSC (https://genome.ucsc.edu/). In addition, using these interfaces, the human DNA sequence of the region of interest was compared with the aligned DNA region of 100 vertebrate species to examine the degree of conservation of the TF RE (hg19 Multiz Alignments of 100 Vertebrates).

### 9. Heat Map Visualization and Hierarchical Cluster Analyses of the TF RE Profile of the UCP genes and the Transcriptomic Data of Human Adipocytes

Hierarchical cluster analyses and heat map visualization were performed with the Morpheus web tool (https://software.broadinstitute.org/morpheus/) using Spearman rank correlation of rows and columns and complete linkage based on the presence or absence of the TF RE in the given gene upstream regulatory region. The same tools were used to visualise relative gene-expression profile of human neck-derived adipocytes (Tóth et al. 2020). RNAseq data is deposited to Sequence Read Archive (SRA) database (https://www.ncbi.nlm.nih.gov/sra) under accession number PRJNA607438. Heatmaps depict the z-scores calculated from the filtered and normalized transcriptomic data of isolated and maintained preadipocytes from subcutaneous and deep neck adipose stromal cells and lipid-laden mature adipocytes differentiated *in vitro* with white and brown protocols described in Tóth et al. (2020).

### 10. Comparison of the Identified Putative TF REs in the Regulatory Sequence of UCP1 with the Corresponding Data Found in the Chip Based Databases

The list of putative TF REs identified by the EPD interface (https://epd.epfl.ch) at p<1e-4 confidence level in human *UCP1* was compared with similar data from public ChipSeq databases. ChipSeq databases collect experimentally validated DNA binding data for TFs to the promoter of individual genes. Two ChipSeq databases, ChiPBase (https://rnasysu.com) and TFlink (https://tflink.net/), were used to identify proteins/TFs binding in the regulatory DNA sequence of the human *UCP1* gene range between -5000 bp and +1000 bp.

### 11. Identification of Repeat Elements in the Genome

The RepeatMasker interface (https://www.repeatmasker.org/) integrated into the UCSC genome browser (https://genome.ucsc.edu/) was used to identify, categorize and determine the exact position of repeat elements in the human *UCP1* promoter (GRCh 37/hg19 by NCBI Gene, Chr4: 141.489.115-141.495.114).

### 12. Literature Mining Using a Custom Script

We programmed a simple Python script which uses the Entrez E-utilities (Sayers 2009) ESearch and EFetch to query the PubMed database and retrieve relevant abstracts. The script summarizes the search results in a convenient, human-readable tabular format which is manually examined by the researcher.

We compiled a list of abbreviations of TFs of interest (e.g. HOXA5, JUN, TEAD1, etc.) based on the JASPAR database; Altogether the list of TFs we examined comprised 579 items.

In addition, we added the UCP keyword to the search.

For each TF in the list, the script iteratively executes searches for each pair consisting of that TF and a keyword, e.g. HOXA5 and UCP1, HIF1A and UCP1 etc. These pairs are concatenated with a ‘+’ character and passed to ESearch, e.g. *‘HOXA5+UCP1’*. When searching for a complex of several TFs rather than a single TF, e.g. EWSR1 and FLI1, the constituent TFs in the complex were also combined with a ‘+’ character in the search, e.g. *‘EWSR1+FLI1+UCP1’*.

When searching for TF abbreviations that are identical or similar to a regular English word, in particular Ar (for which the database returns hits containing the verb form ‘are’), CLOCK (English noun), REST (English noun and verb), MAX (short for maximum and a name) and JUN (very frequent abbreviation of the month June), we found that simply searching for these abbreviations returns a very large number of abstracts which contain the English word rather than the TF in question. Thus we had to add a further search term to disambiguate the abbreviation and narrow down the search to abstracts which contain a reference to the TF that we were interested in. We found that searching for *“Ar androgen receptor”, “CLOCK circadian”, “REST RE1 silencing”, “MAX MYC associated x”,* and *“JUN proto-oncogene”* respectively constrained the search effectively so that the results contained few irrelevant hits.

All combinations of TFs and the keyword were searched, i.e. in total 579 searches were executed. For each such combined TF + keyword search expression, the ESearch tool returns a list of PubMed ids of articles the abstract of which contains both the TF and the keyword. For each TF the number of abstracts containing the keyword is summarized in a table by the script. In addition, the full text of all abstracts is downloaded and saved along with the bibliographical data of the corresponding articles, as well as the URL of the full text of the article where available up to a pre-set maximum number, e.g. 50 abstracts.

Finally, the downloaded abstracts were manually reviewed. We considered the use of advanced natural language processing techniques, in particular large language models, to partially automate the review of the abstracts, but this was judged unnecessary for this specific set of results.

### 13. Identification of TF Motif Activity in Different Tissues and Exploration of TF Interaction Partners Using the ISMARA Online Platform

We obtained RNA-seq expression profiles from 16 human cell types using the Illumina Body Map 2 dataset (GEO accession GSE30611), accessible within the Integrated System for Motif Activity Response Analysis (ISMARA) online platform (https://ismara.unibas.ch/supp/dataset1_IBM_v2/ismara_report/) (Balwierz et al. 2014). We employed ISMARA to analyze the motif activity of putative species-specific and species-common homeodomain family TFs across different human tissues for identification of TF motif activity. By leveraging the ISMARA platform, we investigated the motif activity patterns of these TFs, providing insights into their potential roles in tissue-specific gene regulation. Furthermore, we explored potential TFs and their firs-level TF interaction partners using ISMARA to uncover regulatory networks and complex formations, such as enhanceosomes.

### 14. Statistical Analyses

Statistical analysis was carried out using the stats package in R. Correlation analysis of *UCP1* gene expression with TFs showing gene expression in adipocytes from the cervical region (Tóth et al. 2020) was performed using Sperman’s rank correlation. For multiple comparisons, an adjusted p-value was calculated using Holm’s test, with p < 0.05 considered statistically significant (Suppl. Table 3).

## Supplementary Table Legends

**Supplementary Table 1** The table lists TFs identified through RE analysis in the promoters of human, mouse, and rat UCP1, UCP2, and UCP3 genes, categorized by species and genes, employing the EPD interface (p<1e-4) for identification. Subgroup categorization is based on the Venn diagram representation presented in Figure 1D.

**Supplementary Table 2** The table presents the list of TFs identified through RE analysis in the promoters of human, mouse, and rat UCP1, UCP2, and UCP3 genes. The listing includes the TFs’ relative positions to the TSS, categorized by species and genes, utilizing the EPD interface (p<1e-4, p<1e-5).

**Supplementary Table 3** The table presents correlation coefficients depicting the associations between the gene expression profile of human UCP1 and the TFs identified within human UCP1 promoters using putative REs. The data utilized for this analysis were derived from RNA sequencing of human adipocytes obtained from the cervical region.

**Supplementary Table 4.** The table shows the TF abbreviations and the number of abstracts where they occur together with the keyword UCP1 in publications in the PubMed database.

## DECLARATIONS

### Ethics approval and consent to participate

Not Applicable

### Consent for Publication

Not Applicable

### Availability of data and material

RNAseq data have been deposited to Sequence Read Archive (SRA) database (https://www.ncbi.nlm.nih.gov/sra) under accession number PRJNA607438. We automated the retrieval of EPD data and the screening of abstracts from the PubMed database using custom scripts that were made publicly available on GitHub (https://github.com/DEpt-metagenom/TFRE-in-UCP1-promoter). The main and supplementary figures and tables are available in the Mendeley database: https://data.mendeley.com/preview/z5b6stz3zx?a=828a4650-5239-487a-b17f-43c667b02454

### Authors’ contributions

Conceptualization, BBT., methodology, BBT., GH, EV, GP, and LL; software, BBT., GH, EV, GP and LL; formal analysis, BBT and GP; visualization, BBT and LL; investigation, BBT.; data curation, BBT; writing—original draft preparation, BBT.; writing—review and editing, BBT., PG, LL, LF; project administration, LL;

## Acknowledgements

We extend our appreciation to Professors Jan Nedergaard and Barbara Canon for their insightful feedback and valuable suggestions to enhance the clarity of the manuscript. On behalf of the project "Bioinformatic analysis of third generation sequencing data" ("Harmadik generációs szekvenálási adatok bioinformatikai elemzése") we are grateful for the possibility to use HUN-REN Cloud (see Héder et al. 2022; https://science-cloud.hu/) which helped us achieve the results published in this paper.

## SUPPLEMENTARY FIGURES

**Supplementary Figure 1.**
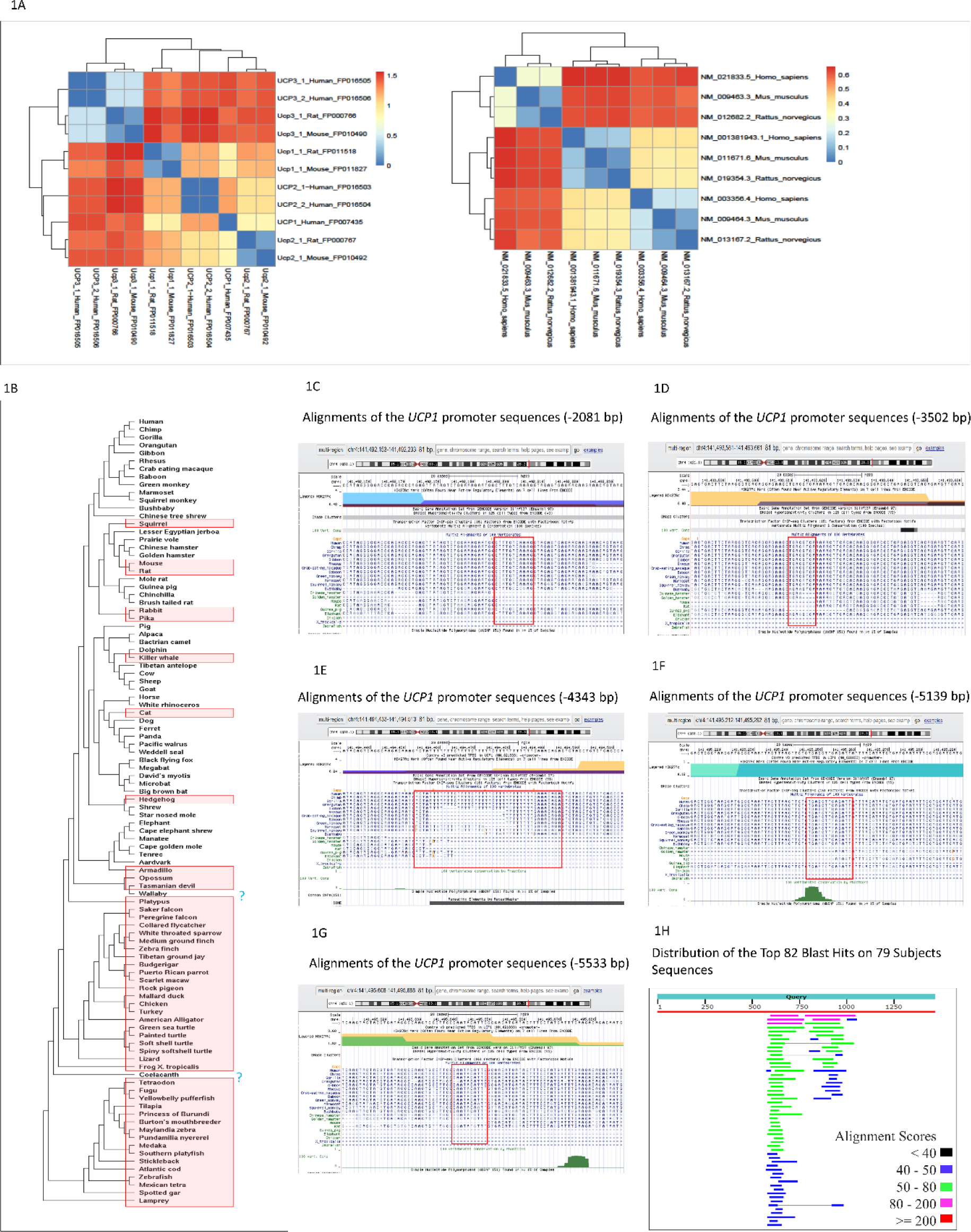
Heatmaps show the pairwise genetic distances of the nucleotid sequences of the *UCP1*, *UCP2* and *UCP3* genes’ regulatory and coding regions in humans, mice and rats (pheatmap program in R environment). **1B** The phylogenetic tree shows the presence and the absence (red box) of the human UCP1 promoter DNA fragment spans between -2083 bp and -3502 bp (GRCh37/hg19: Chr4 141,492,160-141,493,617) in *UCP1* promoter of different vertebrate species according to the Multiz Alignment of 100 Vertebrates database in the UCSC genome browser. No information is available for the three genera marked with blue question marks. **1C – 1G** Alignments of the *UCP1* promoter sequences of different species highlight the palindromic nucleotide sequences of the beginning and end of the DNA fragments (1419 bp and 658 bp) missing from the mouse and rat *UCP1* promoter according to the Multiz Alignment of 100 Vertebrates database in the UCSC genome browser. **1H** Graphical summary showing the human genomic occurrence of the *UCP1* promoter DNA fragment (1419 bp) absent from the mice and rat *UCP1* promoter (https://blast.ncbi.nlm.nih.gov/). The graphic is an overview of the database sequences aligned to the query sequence. These are represented by horizontal bars colored coded by score and showing the extent of the alignment on the query sequence. Separate aligned regions on the same database sequence are connected by a thin grey line.

**Supplementary Figure 2.**
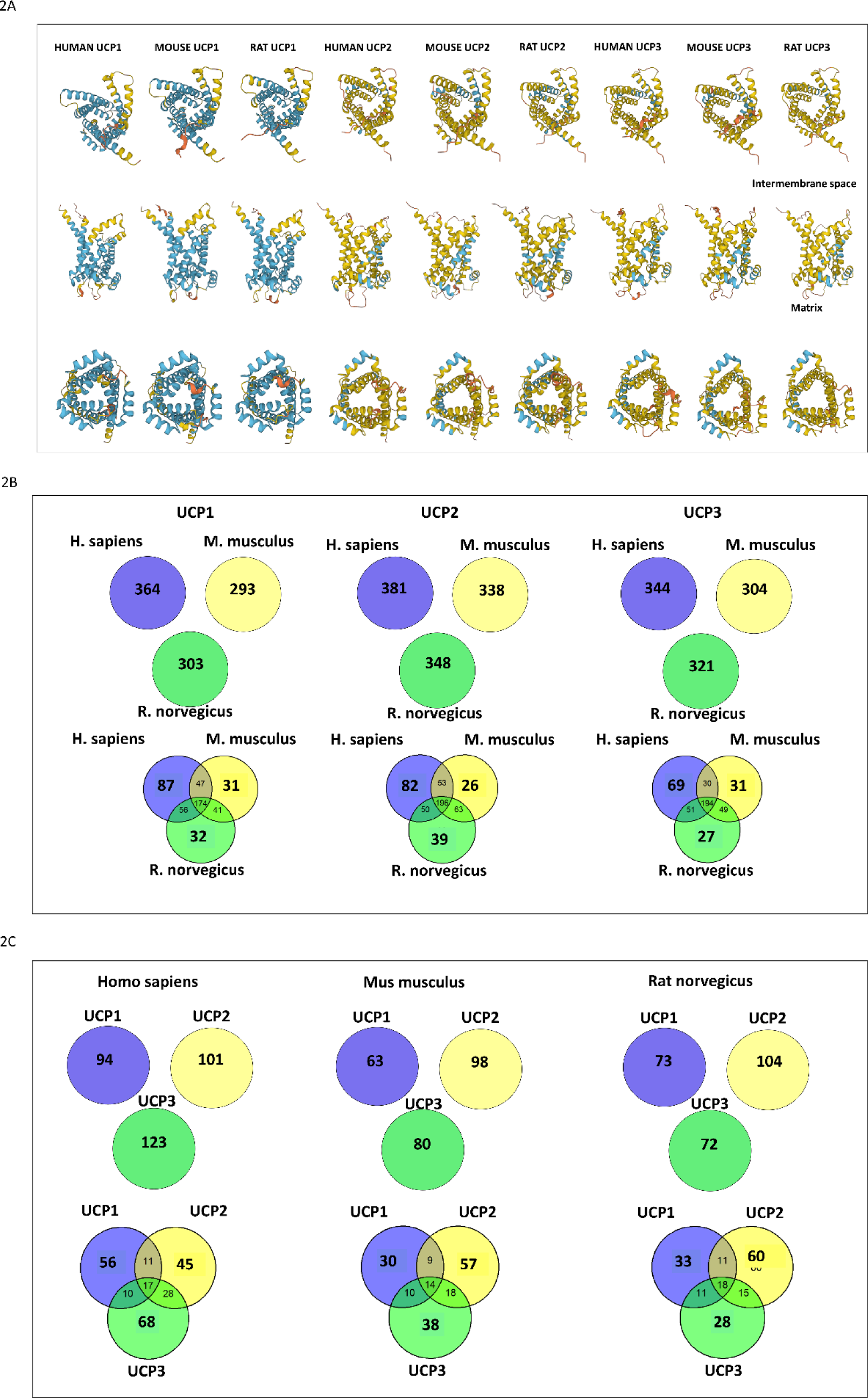
**2A** The spatial structure of human, mouse and rat *UCP1*, *UCP2* and *UCP3* proteins predicted by AlphaFold. The figures show the predicted spatial structure of the protein from the inter-membrane face, the membrane-embedded side and the matrix face. The blue color indicates that for the *UCP1* gene, the model predicts the spatial structure with high confidence for most of the part of the protein for all three species, while for the other UCPs, the model confidence value is generally low. Typically, a short highly disordered portion facing the matrix is well conserved in the UCP proteins of all three species. This potentially context-dependent variable spatial structure may affect the function of the protein at a given time. **2B** Venn diagram showing the number of TF REs identified in the promoter (-5000 bp –TSS – +1000 bp) of human, mouse and rat *UCP1*, *UCP2* and *UCP3* genes using the EPD interface, with probability level p<1e-4. Merged Venn diagrams among species show the number of unique and common TF REs in *UCP1*, *UCP2* and *UCP3* genes. **2C** Venn diagram showing the number of TF REs identified in the promoter (-5000 bp –TSS – +1000 bp) of human, mouse and rat *UCP1*, *UCP2* and *UCP3* genes using the EPD interface, with probability level p<1e-5. Merged Venn diagrams among *UCP* genes show the number of unique and common TF REs in human, mouse and rat *UCP* genes (Venn program in R environment).

**Supplementary Figure 3.**
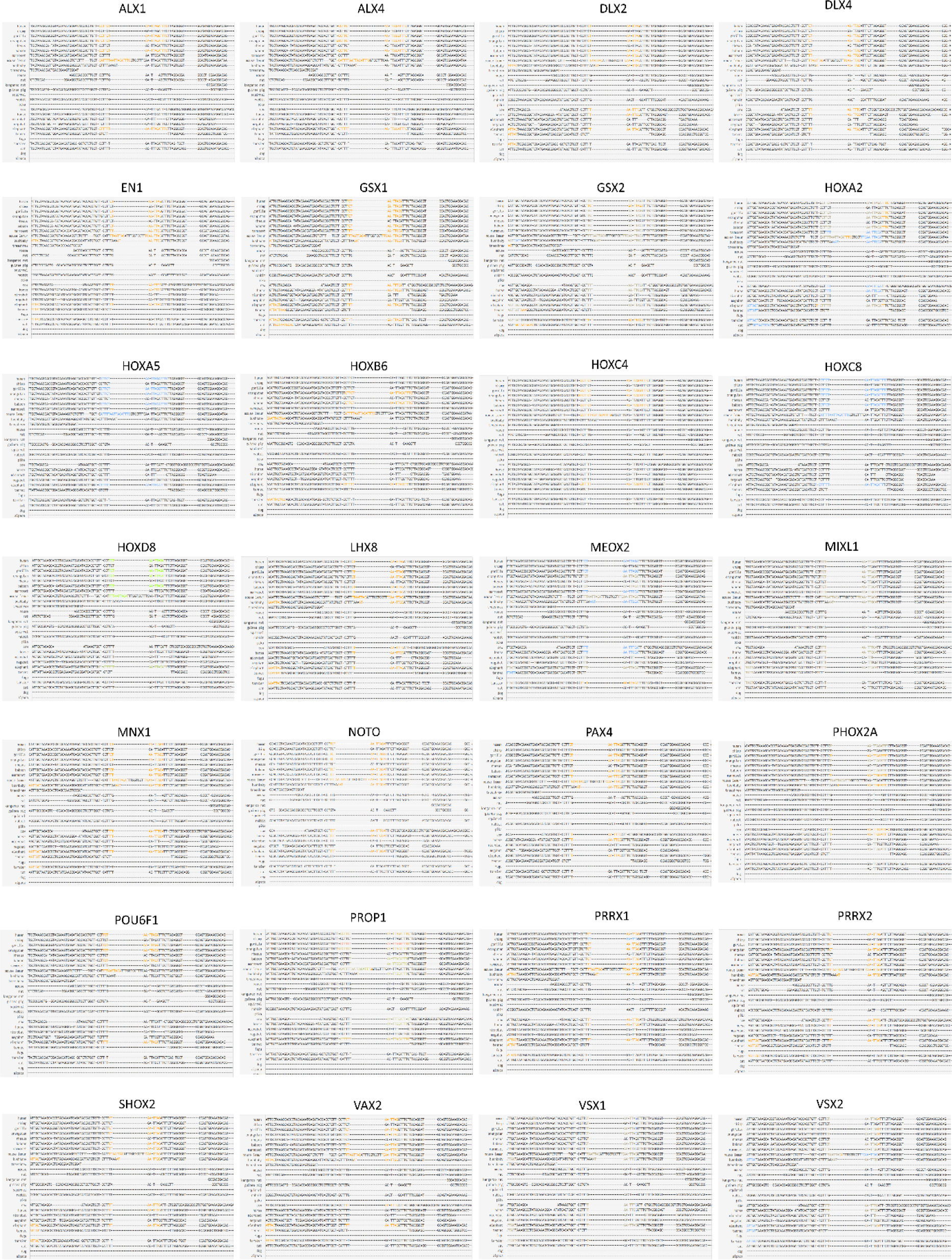
In the aligned *UCP1* promoter sequence of different species at position -690 bp, we investigated the presence of TF binding sequences on the ConTra v3 platform, which gave positive search results on the EPD interface, as well as those that were not tested using EPD but have a potential RE based on their TF class classification. Stringency: core = 0.95, similarity matrix = 0.85.

**Supplementary Figure 4.**
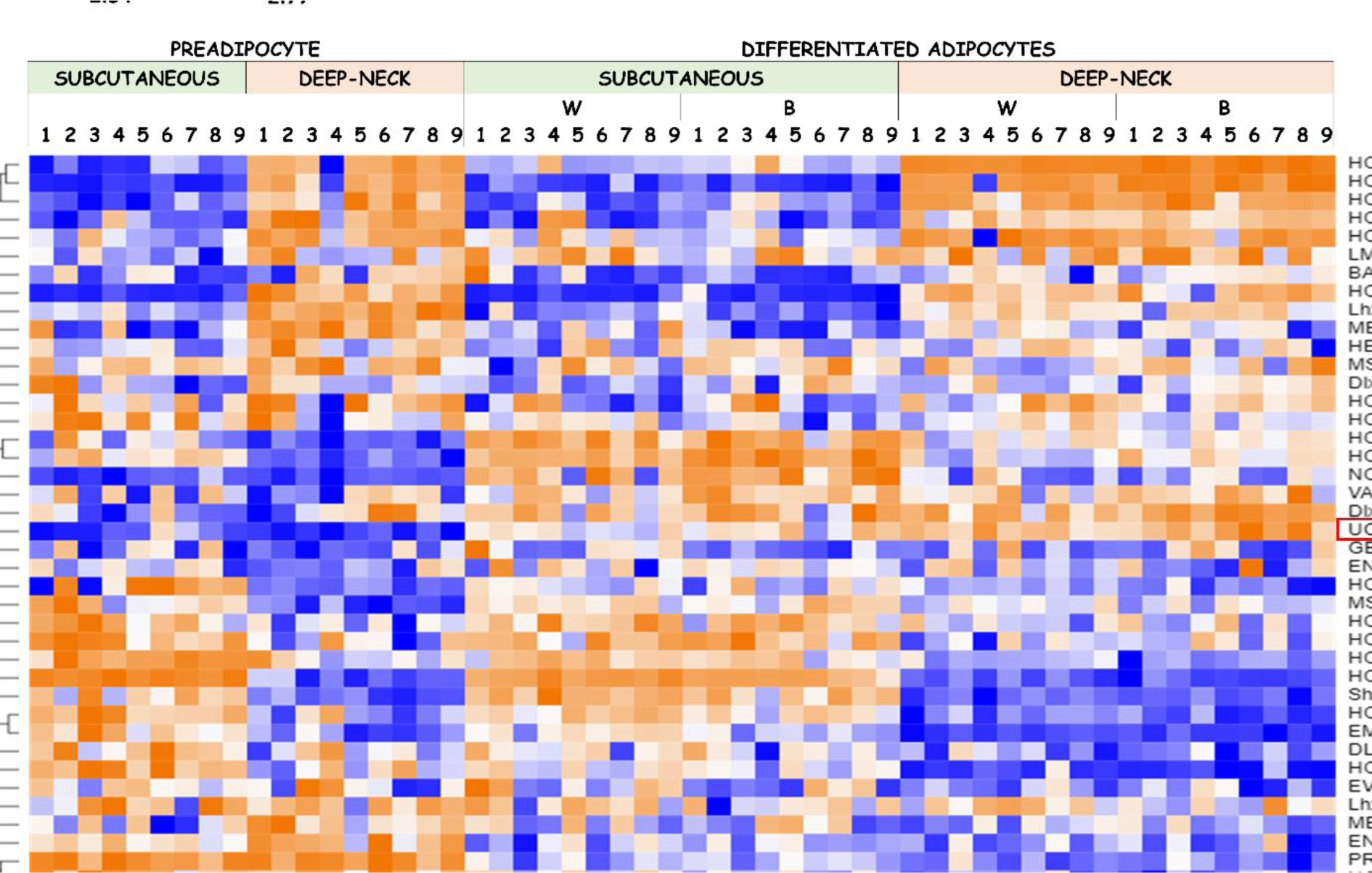
The heatmap depicts the relative gene expression profile of homeodomain Factor 1 class TFs (based on the JASPAR database) in adipocytes of cervical origin. For comparison, *UCP1* is also shown (Morpheus interface). Hierarchical cluster analysis based on the Spearman Rank similarity matrix identifies TFs with gene expression profiles similar to *UCP1*. Adipocytes were differentiated using a white (W) or brown (B) differentiation protocol.

**Supplementary Figure 5.**
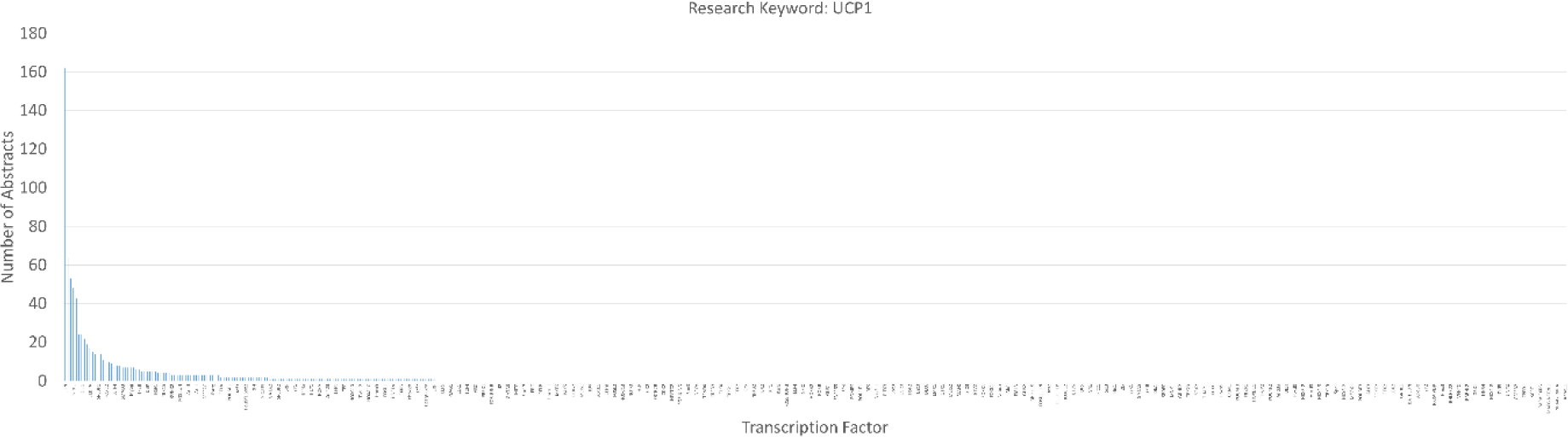
The bar graphs show the number of abstracts deposited in the PubMed database in which the TF under study appears together with the keyword UCP1, highlighting the “longtail” distribution of the data. On the x-axis only a few representative TFs are named, the y-axis shows the number of abstracts (Excel).

**Supplementary Figure 6.**
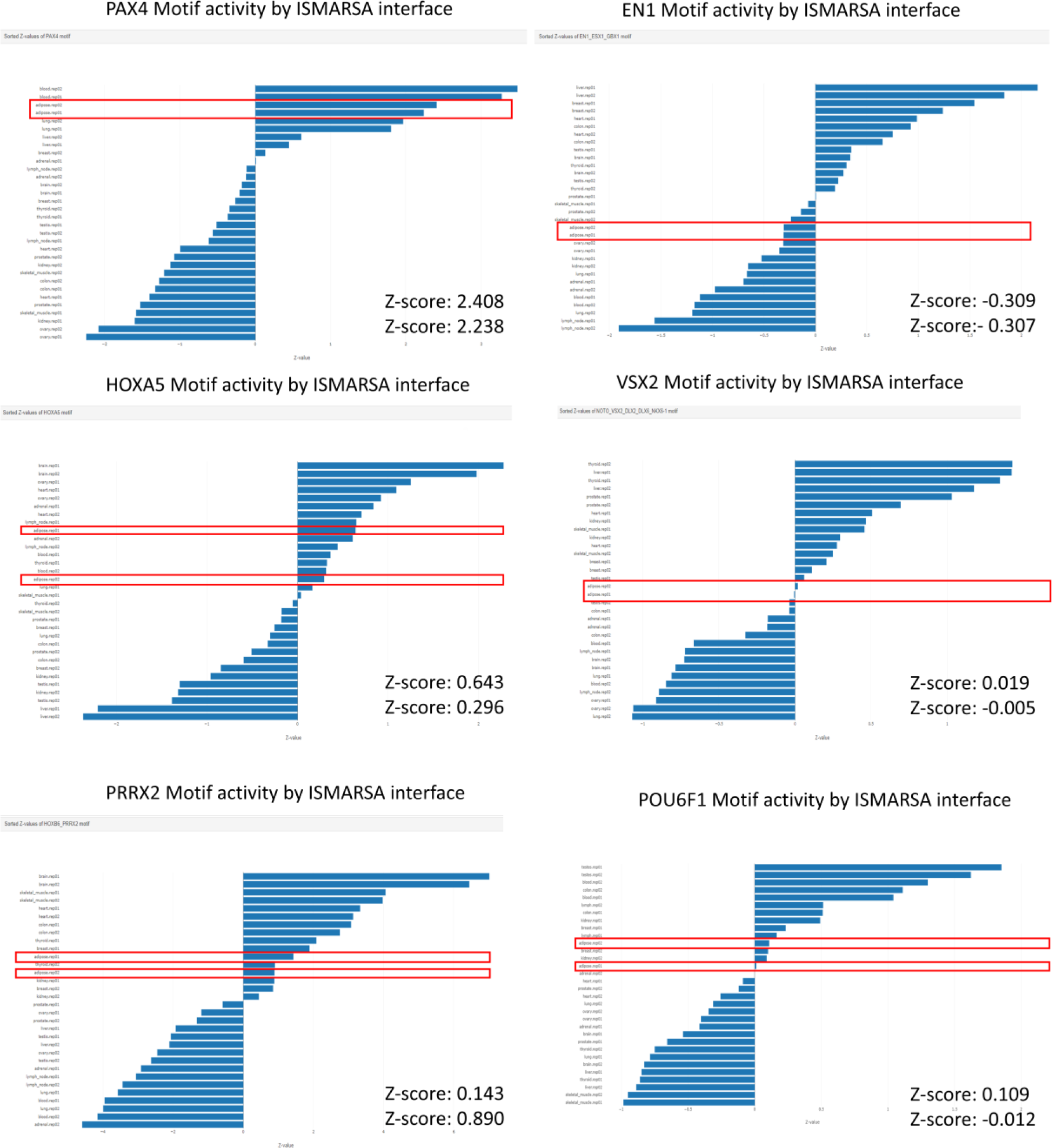
Bar graphs show the relative activity (z-score) of the putative species-specific and species-common homeodomain family TFs under study assessed per human tissue (according to ISMARA interface). The red box highlights the result recorded in adipose tissue samples.

**Supplementary Figure 7.**
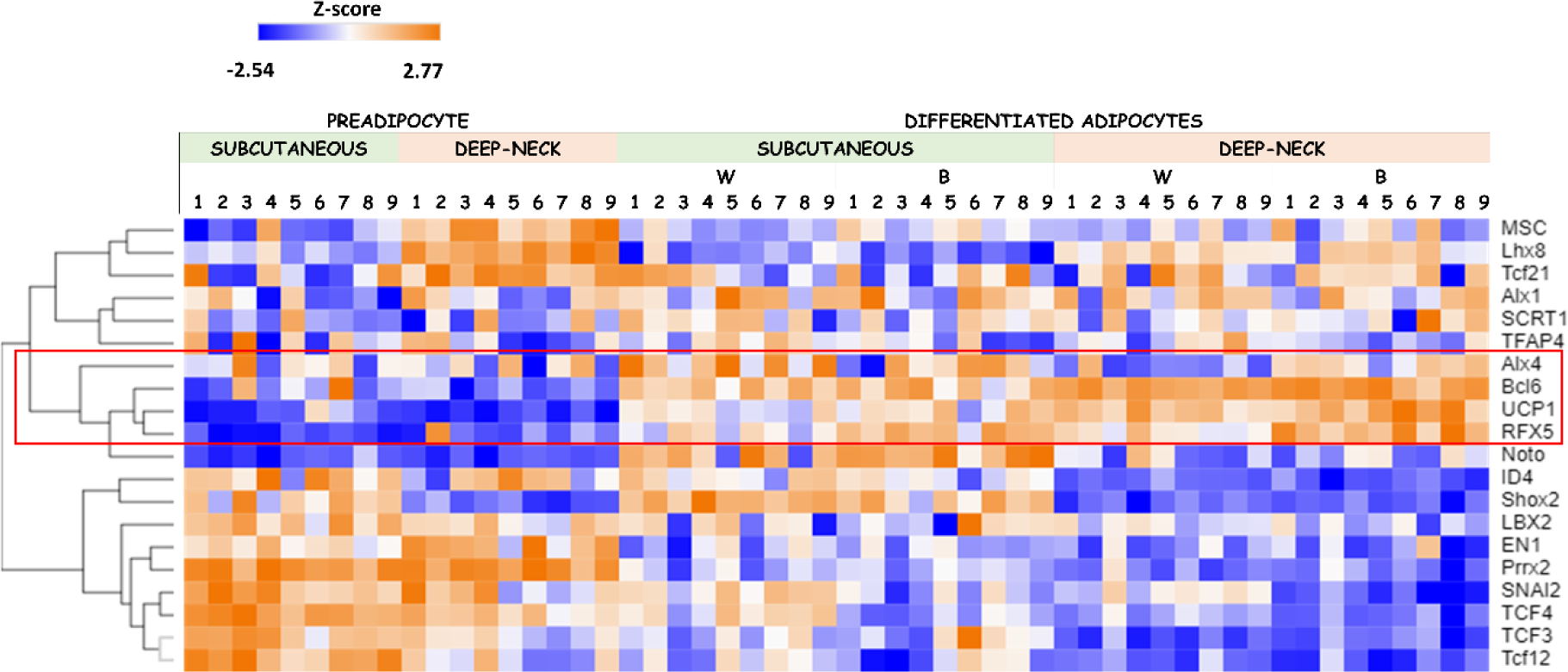
The heatmap shows the relative gene expression profiles of TFs, for which a binding sequence was identified at -690 bp and near +/- 40 bp in the promoter of human *UCP1* using EPD interface (p<1e- 4). For comparison, *UCP1* is also shown. Hierarchical cluster analysis based on Spearman Rank similarity matrix identifies TFs with gene expression profiles similar to *UCP1* (red box) (Morpheus interface). Adipocytes were differentiated using a white (W) or brown (B) differentiation protocol.

**Supplementary Figure 8.**
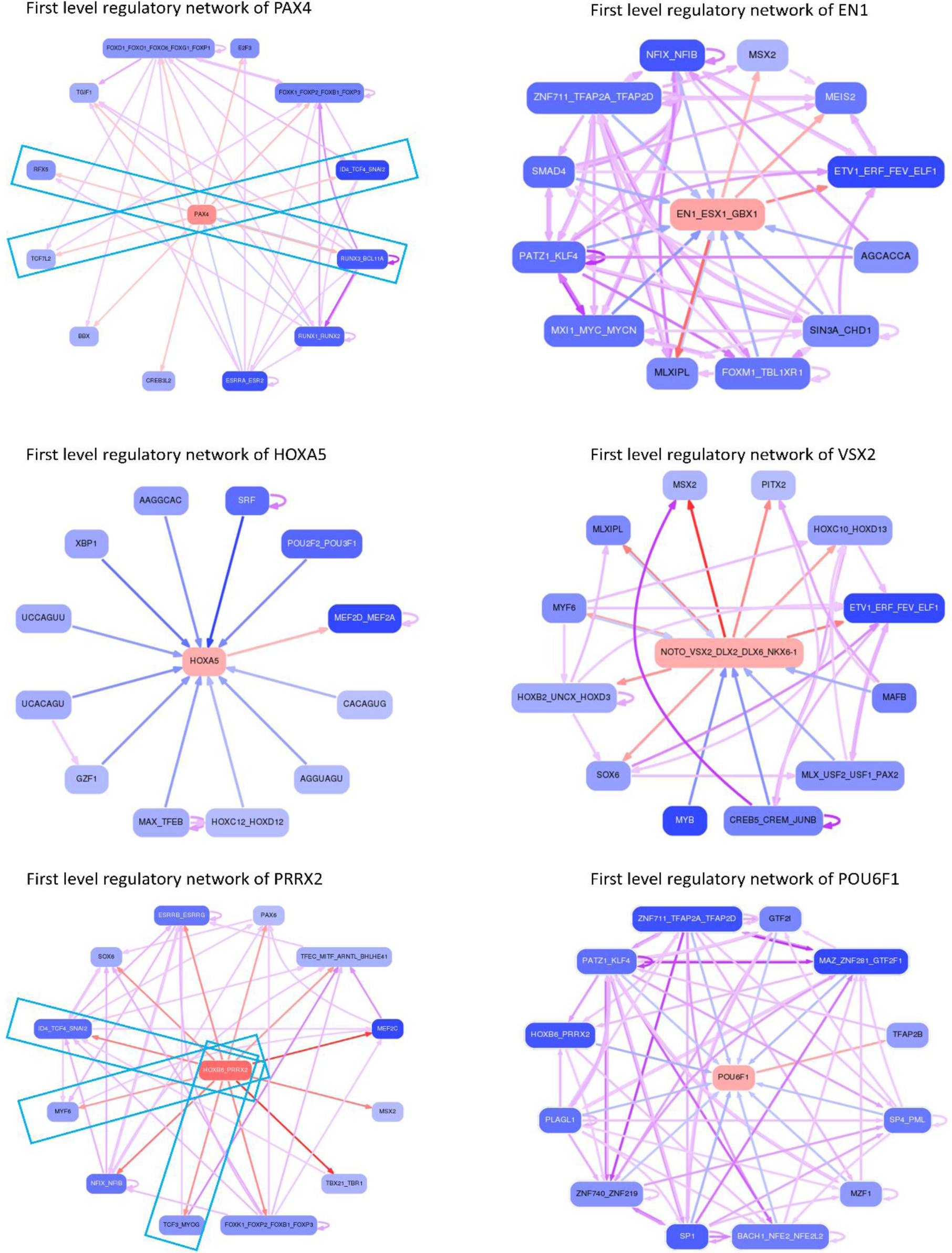
Interactomes show the first level of the regulatory network of the putative species- specific and species-common homeodomain family TFs under study, such as PAX4, EN1, HOXA5, DLX2, PRRX2 and POU6F1 according to the ISMARA interface. The blue box highlights the partners that were recognized as having a RE close to (within +/- 40 bp of) the RE of the TF of interest.

**Supplementary Figure 9.**
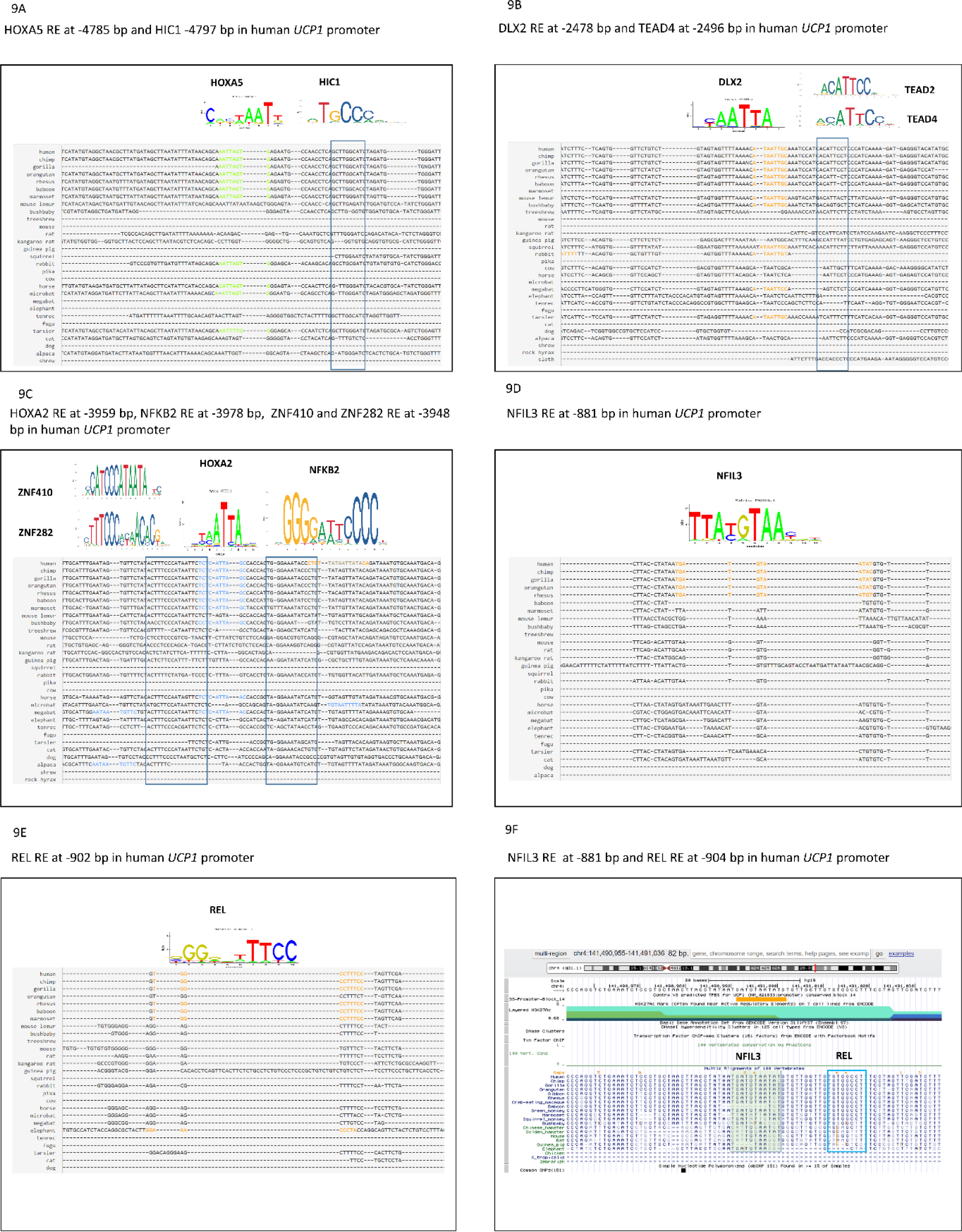
Alignment of *UCP1* promoter sequences from different species at different positions on the ConTra v3 platform. Human *UCP1*-specific TF REs are highlighted in different colors, and nearby TF REs are outlined in a blue box. **9A** shows the presence of the HOXA5 TF RE, unique to the human *UCP1* promoter at -4785 bp, and the nearby HIC1 RE at -4797 (EPD interface data). **9B** shows the presence of the DLX2 TF RE, unique to the human *UCP1* promoter at -2478 bp, and the nearby TEAD1-4 RE at -2496 (EPD interface data). **9C** shows the presence of the HOXA2 TF RE, unique to the human *UCP1* promoter at -3959 bp, and the nearby NFKB RE at -3978, and ZNF410 and ZNF282 at - 3948 bp (EPD interface data). **9D** shows the presence of the NFIL3 TF RE, unique to the human *UCP1* promoter at -881 bp, and **9E** the nearby REL RE at -902 bp (EPD interface data). **9F** shows the presence of the NFIL3 TF RE at -881 bp and the nearby REL RE at -902 bp across species in UCSC genome browser (ConTra v3 and EPD interface data).

**Supplementary Figure 10.**
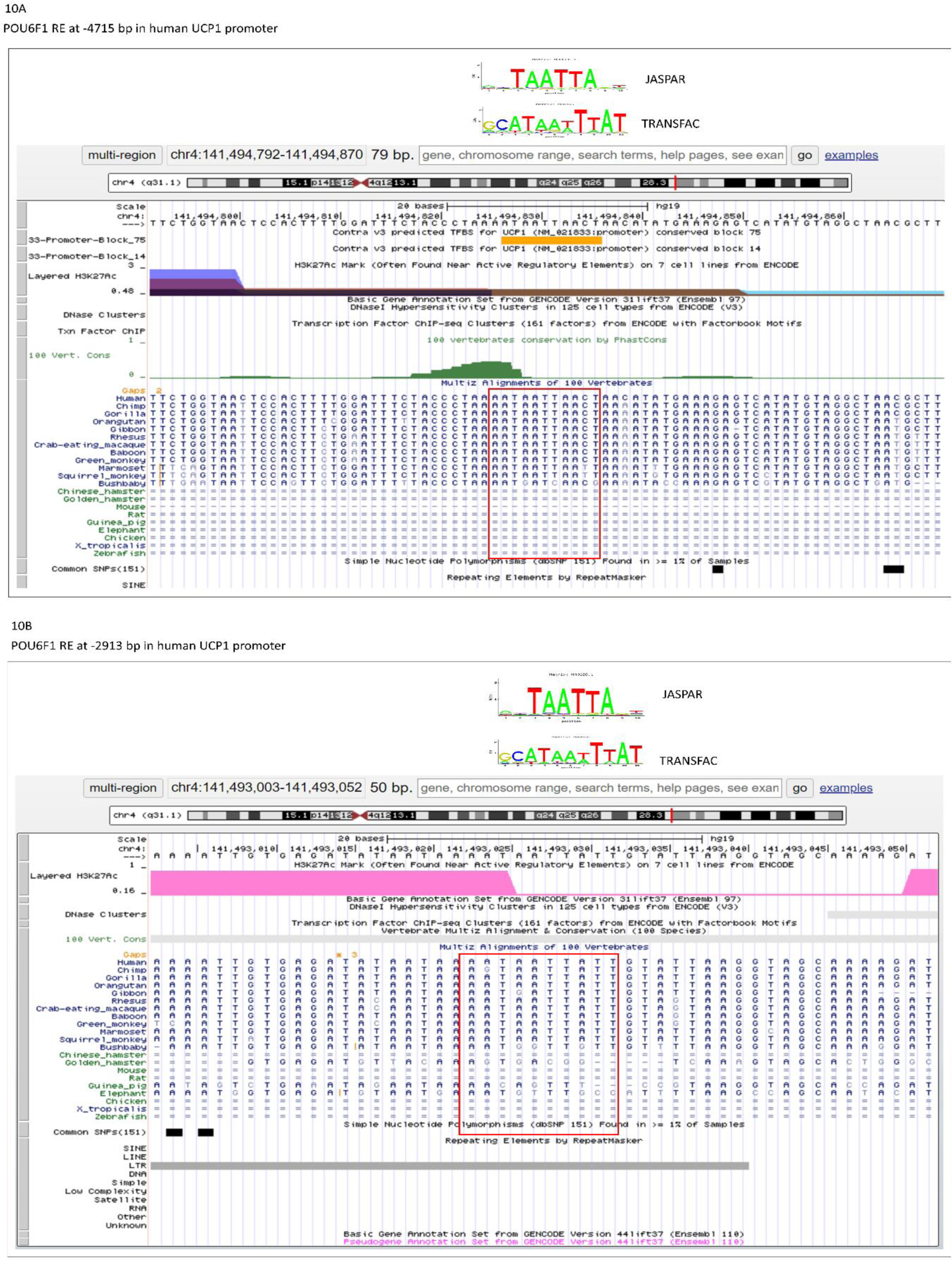
Alignment of the *UCP1* promoter sequences of different species **10A** at -4715 bp position and **10B** at -2913 bp position to investigate the presence of POU6F1 TF-binding sequences in human, mouse and rat promoter according to the Multiz Alignment of 100 Vertebrates database in the UCSC genome browser.

**Supplementary Figure 11.**
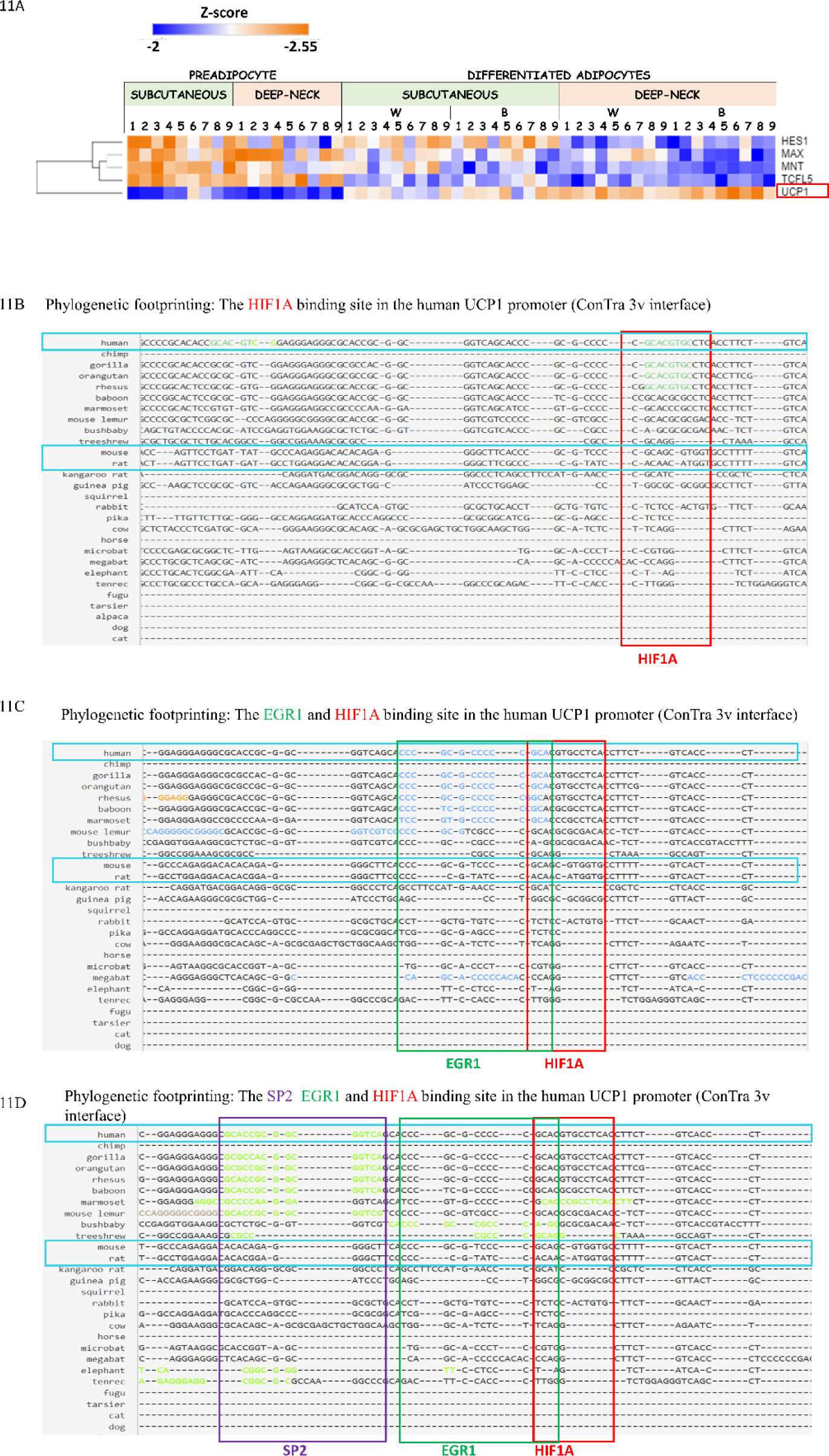
**11A** Heatmap showing the relative gene expression profile of common TFs identified in the promoters of human *UCP1*, *UCP2* and *UCP3* around -264 bp missing the binding sequence in the promoters of mouse and rat orthologous genes using EPD interface (p<1e-4) in cervical derived adipocyte samples. For comparison, *UCP1* is also shown. Hierarchical cluster analysis based on the Spearman Rank similarity matrix identifies TFs with similar gene expression profiles to *UCP1*. (Morpheus Interface) **11B** Phylogenetic footprinting: the green color/red box highlights the HIF1A binding site in the *UCP1* promoter at -264 (ConTra v3 interface). **11C** Phylogenetic footprinting: Blue color/green box highlights the EGR1 RE at -257 and and red box the HIF1A RE at -264 bp in the *UCP1* promoter (ConTra v3 interface). **11D** Phylogenetic footprinting: the green color/purple box highlights the SP2 RE at -223 bp, the green box shows the EGR1 RE at -257 bp and the red box the HIF1A RE at -264 bp in the *UCP1* promoter (ConTra v3 interface). Adipocytes were differentiated using a white (W) or brown (B) differentiation protocol.

